# Therapeutic Interruption of T Cell Development Generates High-Affinity T Cells That Escape Exhaustion and Improve Cancer Immunotherapy

**DOI:** 10.1101/2022.01.19.476935

**Authors:** Erica Dhuey, Olivia Oldridge, Roshan Ravishankar, Hannah Dada, Yongjun Yu, Mark S. Anderson, Andy J. Minn

## Abstract

Availability of effective anti-tumor T cells is limited by cancer immunoediting, which depletes neoantigens, and central tolerance, which eliminates developing T cells with high-affinity T cell receptors (TCRs) against tumor self-antigens. Remaining tumor-reactive T cells are often exhausted after immune checkpoint blockade (ICB). Whether endogenous T cells with high-affinity TCRs against tumor self-antigens can be generated to circumvent exhaustion and reject neoantigen-poor tumors is unclear. We show that transiently interrupting central tolerance through RANKL blockade unleashes T cells possessing TCRs with self-reactive features that enable ICB to reject poorly immunogenic tumors. Upon recognition of tumor self-antigens, these T cells exhibit enhanced TCR signaling, enrichment in NFAT/AP-1 genes, and lymph node priming. Consequently, memory-precursor T cells against tumor self-antigens are generated, avoid exhaustion, and become effector-memory cells with transcriptional features associated with clinical ICB response. Thus, interrupting central tolerance provides T cells with tumor-directed autoreactivity that avoid exhaustion and improve immunotherapy.

## INTRODUCTION

Immune checkpoint blockade (ICB) has shown tremendous clinical promise, particularly against aggressive and advanced tumors. Blocking antibodies against immune checkpoints CTLA4 and PD1 have delivered impressive responses against several solid tumors, including melanoma, bladder cancer, and non-small cell lung cancer (Ribas and Wolchok, 2018). However, most patients do not respond and/or relapse after ICB therapy. Furthermore, many of the most common and deadly cancers, such as breast and pancreatic cancers, are typically poorly responsive. Thus, developing therapies that overcome major barriers to ICB response hold substantial promise.

One critical requirement for ICB response is tumor immunogenicity, which is the ability of tumors to elicit T cell responses. Tumor immunogenicity is strongly impacted by the quality and quantity of tumor epitopes displayed by major histocompatibility complex (MHC) molecules on the cell surface for T cells to survey (Gubin et al., 2014; Rizvi et al., 2015; Schumacher and Schreiber, 2015). One important class of tumor-specific epitopes are neoantigens, which are novel peptides derived from somatic mutations. The number of somatic mutations (tumor mutational burden) positively correlates with both neoantigen load and with ICB response, and may account for up to 55% of the difference in response rates between cancer subtypes (Jardim et al., 2021). However, many patients have a paucity of neoantigens, resulting in poorly antigenic tumors that do not benefit from ICB therapy (Schumacher and Schreiber, 2015). Even among tumors with an initially favorable repertoire of neoantigens, the tumor clones with highly immunogenic neoantigens are preferentially eliminated by the immune system and contribute to treatment relapse. Thus, interventions that can increase tumor immunogenicity despite a dearth of strong neoantigens may be required to augment ICB therapy for most cancer patients.

Several therapeutic approaches have been developed to target tumors with poor neoantigen availability. These include adoptive cell therapies (Lim and June, 2017) and cancer vaccines against tumor self-antigens (Farkas and Finn, 2010), which are antigens found on normal host tissue but are often aberrantly expressed by cancer cells and exposed to immune recognition. However, since tumor self-antigens are germline-encoded, T cells capable of strongly recognizing such antigens are largely eliminated from the repertoire during T cell development in the thymus. In a process known as central tolerance, self-antigens are presented to developing T cells by medullary thymic epithelial cells (mTECs), and the T cells with T cell receptors (TCRs) that bind with high affinity to these self-antigens are deleted (Hogquist et al., 2005). Consequently, high-affinity T cells against self-antigens do not typically accumulate in the periphery, which is critical for preventing autoimmunity. In fact, mice and humans deficient in *Aire*, a transcription factor responsible for expression of self-antigens by mTECs, develop multiorgan autoimmune disease (Anderson et al., 2002; Consortium, 1997; Nagamine et al., 1997). However, in the context of cancer, in which tumors arise from self-tissues, central tolerance severely limits the ability of self-antigens expressed by tumors to elicit T cell responses (Bakhru et al., 2017; Trager et al., 2012; Zhu et al., 2013). Thus, therapies such as ICB or cancer vaccination that rely on generating robust T cells responses against tumor self-antigens are often ineffective (Ryan et al., 2010). Interestingly, mTECs can be temporarily depleted through treatment with a blocking antibody against receptor activator of nuclear factor kappa-B ligand (RANKL) (Khan et al., 2014; Metzger et al., 2013), which through a different mechanism is coincidentally used in the clinical treatment of osteoporosis (denosumab). In fact, RANKL blockade in mice can interfere with the deletion of transgenic T cells that are normally susceptible to central tolerance (Khan *et al*., 2014). These observations suggest that therapies that interfere with T cell development could generate high-affinity T cells against tumor self-antigens that can potentially improve immunotherapy.

Besides insufficient immunogenicity, ICB efficacy is also restricted by T cell exhaustion, a terminal differentiation state characterized by expression of multiple inhibitory receptors, key exhaustion-related transcription factors, and T cell dysfunction. Once T cells commit to terminal exhaustion, often due to chronic antigen stimulation, the effectiveness of immunotherapies are likely severely limited due to epigenetic changes that lock-in the exhausted state (Abdel-Hakeem et al., 2021; Pauken et al., 2016). Recent studies examining human melanoma patients reveal that nearly all tumor-reactive T cells against neoantigens are exhausted (Oliveira et al., 2021). Similarly, T cells against tumor self-antigens are also exhausted and possess low-affinity TCRs, consistent with the effect of central tolerance. Whether T cells with high-affinity TCRs against tumor self-antigens can escape exhaustion, adopt more favorable effector-memory features, and enable ICB to drive better anti-tumor immunity are unclear.

In this study, we use an anti-RANKL antibody to transiently interfere with T cell development in the thymus. Tumor response to ICB and the fate of ensuing endogenous T cells are then linked to the affinity and self-reactive features of their TCRs against a dominant tumor self-antigen. Our work highlights how agents that change the composition of the T cell repertoire can give rise to tumor-directed autoreactive T cells that are resistant to T cell exhaustion and improve immunotherapy response against poorly immunogenic tumors.

## RESULTS

### RANKL blockade improves anti-tumor immunity against poorly immunogenic tumors

We hypothesized that RANKL blockade can transiently disrupt central tolerance to release normally deleted high-affinity T cells against tumor self-antigens that can target poorly immunogenic tumors. We used Res 499 murine melanoma as a model for a poorly immunogenic tumor based on two criteria. First, Res 499 cells are derived from a B16-F10 melanoma tumor that relapsed after ICB-based therapy and have therefore acquired resistance (Twyman-Saint Victor et al., 2015). Second, previous evidence suggests that Res 499 cells have a depletion of predicted neoantigens compared to parental B16 cells (Benci et al., 2019). To confirm and extend these latter findings we compared the landscape of predicted neoantigens from B16 and Res 499 cells using stricter criteria than previously used. Since strong clonal neoantigens are considered more critical to ICB response, we only examined neoantigens with predicted high-affinity (< 100 nM), an allelic frequency consistent with a near- heterozygous genotype, and expression at the RNA level. This revealed that several predicted high-affinity clonal peptide neoantigen are significantly depleted in Res 499 cells compared to B16 cells as determined by a reduction in allelic frequencies to near zero (Figure 1A, lower right quadrant). This includes peptides encoded by *Zic2*, *Lrrc28*, and *Pnp*, which recently were independently identified as top predicted neoantigens in B16 cells (Wert-Carvajal et al., 2021). Moreover, the cumulative distribution frequency of allelic frequencies for all predicted top neoantigens in B16 cells is significantly shifted to lower values in Res 499 cells, consistent with a varied but broad degree of immunoediting (Figure 1B). Thus, these data suggest that Res 499 is a poorly immunogenic tumor as defined by acquisition of ICB resistance and apparent depletion in strong clonal neoantigens.

**Figure 1.**
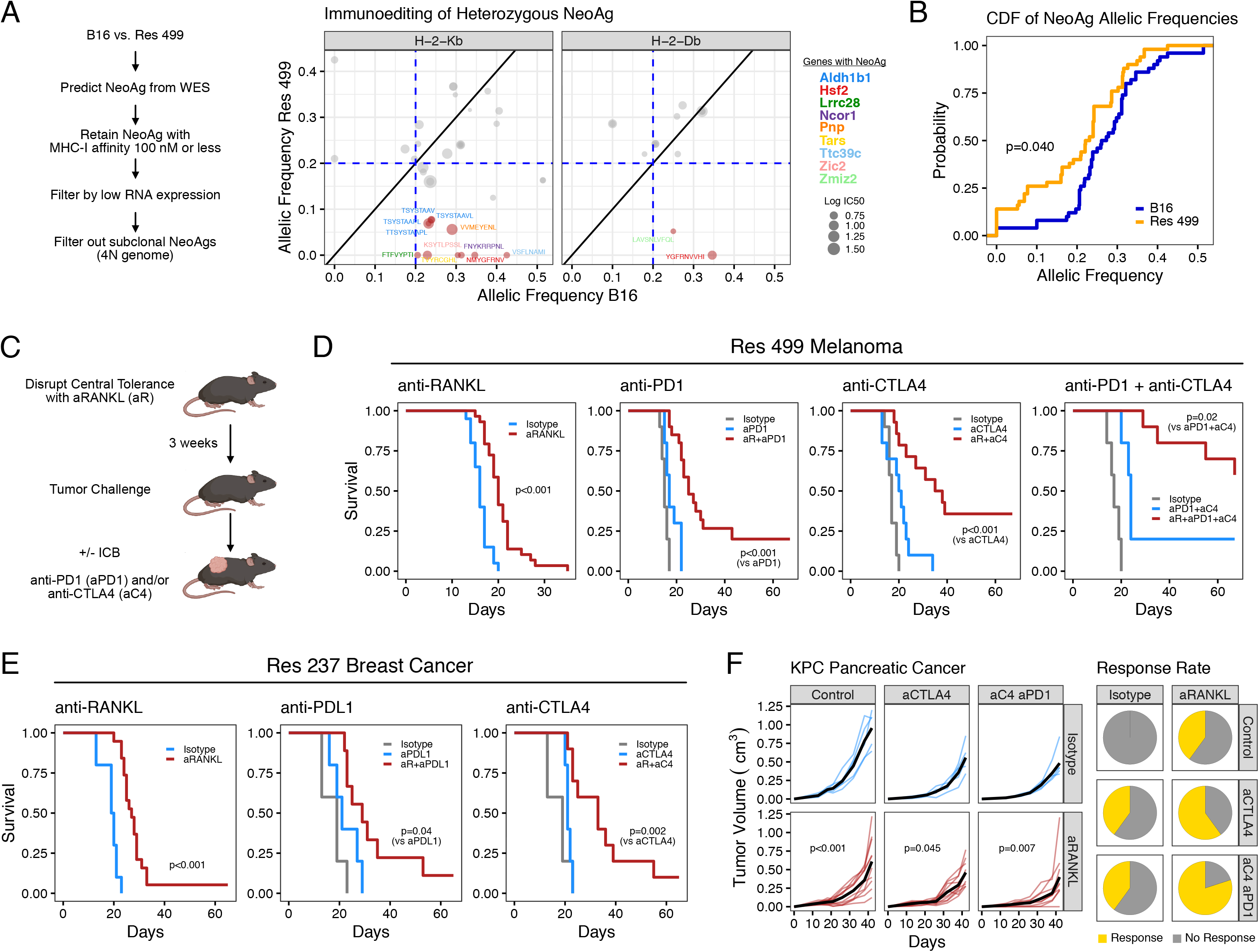
RANKL blockade re-sensitizes neoantigen-depleted ICB-resistant tumors to immune checkpoint blockade. **(A)** Allelic frequencies of predicted neoantigens in B16 and ICB-relapsed Res 499 cancer cells. Peptide antigens for H-2K^b^ and H-2D^b^ are separately shown. Size of dots indicate predicted MHC-I affinity and color indicates gene that encodes the neoantigen peptide. The analysis pipeline is summarized on the left. **(B)** Plot of the cumulative distribution function for the allelic frequencies from (A). P-value is from a KS test. **(C)** Schema of anti-RANKL pre-conditioning 3 weeks prior to tumor challenge with or without subsequent ICB treatment with anti-PD1/PDL1 and/or anti-CTLA4. **(D)** Survival of mice bearing Res 499 tumors after pre-treatment with anti-RANKL or isotype control (n=7-15) or anti-RANKL followed by anti-PD1 (n=5-20), anti-CTLA-4 (n=10-14), or anti- PD1 and anti-CTLA-4 (n=5-10). **(E)** Survival of mice bearing Res 237 ICB-relapsed breast cancer tumors after pre-treatment with anti-RANKL (n=5-15) or after anti-RANKL followed by anti-PDL1 (n=5-9) or anti-CTLA-4 (n=5-10). **(F)** Tumor growth (left) and response rate (right, yellow) of mice bearing Panc 4662 tumors treated with anti-CTLA4 +/- anti-PD1 with or without anti-RANKL pre-treatment (n=5-10). P-values for survival are by log-rank test. Mixed effect model is used for tumor growth analysis. See also Figure S1.

To test how interruption in central tolerance impacts the growth of Res 499 tumors, we administered a blocking antibody against RANKL (anti-RANKL) three weeks prior to tumor inoculation to deplete mTECs (Figure 1C), which was confirmed by monitoring changes in thymocyte composition (Figure S1A-B) as previously reported (Khan *et al*., 2014). Mice with a T cell repertoire pre-conditioned with anti-RANKL demonstrate significantly delayed tumor growth and improved survival compared to isotype control (Figure 1D). Addition of anti-PD1, anti-CTLA-4, or both after anti-RANKL further improves survival and leads to durable therapy response. All mice that achieved a complete response were then re-challenged and none of the mice treated with dual ICB therapy experienced tumor outgrowth. Thus, RANKL blockade enhances anti-tumor immunity and improves ICB response against poorly immunogenic tumors. To extend the effect of anti-RANKL to other poorly immunogenic tumors, we performed analogous experiments with Res 237 breast cancer cells, which is an ICB-relapsed cell line derived from TS/A breast cancer (Twyman-Saint Victor *et al*., 2015), and Panc 4662 pancreatic cancer cells derived from neoantigen-poor tumors arising from *Kras^G12D^*:*Trp53^R172H^*:*Pdx-1*-*Cre* mice (Bayne et al., 2012). In both tumor models, anti-RANKL prolonged survival and improved response to CTLA4 and PD-1/PDL1 blockade (Figure 1E-F). These results further demonstrate that RANKL blockade improves ICB response against multiple poorly immunogenic cancer types.

### CD8 T cells mobilized after RANKL blockade are required to improve anti-tumor immunity

To evaluate whether the effect of RANKL blockade is T cell-dependent, we utilized RAG1 knock-out mice implanted with Res 499 melanoma. This revealed that the delayed tumor growth resulting from anti-RANKL is ablated in the absence of adaptive immunity (Figure 2A). Moreover, elimination of CD8 T cell using depleting antibodies results in a similar effect, indicating that CD8 T cells are specifically required (Figure 2B and S2A). In contrast, depletion of CD4 T cell and NK cell have no significant effect on the efficacy of anti-RANKL (Figure 2C-D and S2A). Together, these data indicate that CD8 T cells and not NK or CD4 T cells are required to inhibit tumor growth in response to RANKL blockade.

**Figure 2.**
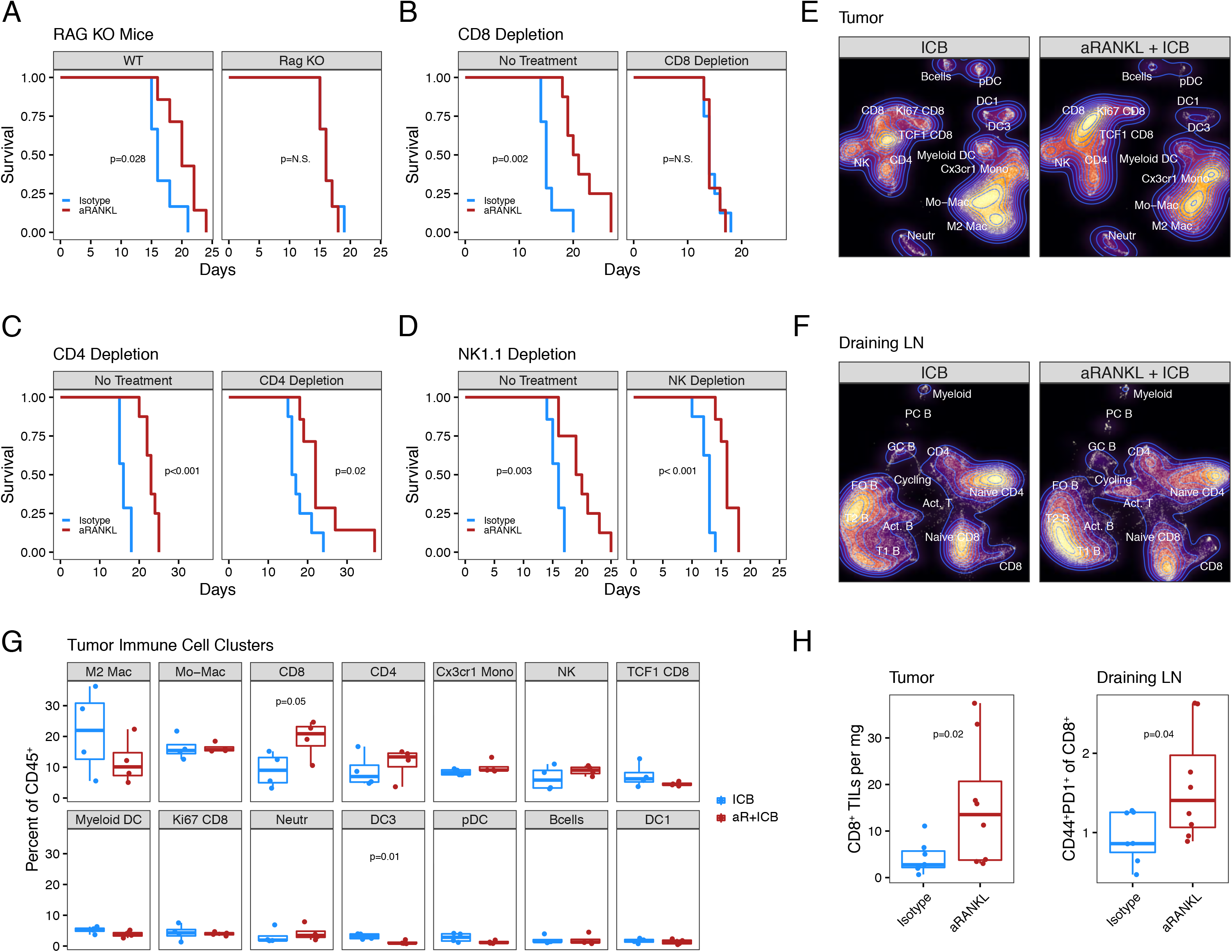
RANKL blockade mobilizes CD8 T cells to delay tumor growth and prolong survival. **(A)** Survival of wildtype (WT) or RAG knockout (KO) mice bearing Res 499 tumors and pre- treated with anti-RANKL or isotype control (n=6-7). **(B-D)** Survival of mice bearing Res 499 tumors after pre-treated with anti-RANKL and antibody- mediated depletion of CD8 T cells (n=7-8) **(B)** CD4 T cells (n=7-8) **(C)** or NK1.1-positive cells (n=7-8) **(D)**. **(E-F)** Frequency densities of immune cells from scRNA-seq clustered and projected on a UMAP plot. Data are pooled replicates (n=4) of Res 499 tumors **(E)** or draining lymph nodes **(F)** harvested at day 15 from mice treated with anti-CTLA4 plus anti-PD1 (ICB) with or without anti- RANKL pre-conditioning (aR). **(G)** Percentage of CD45^+^ cells in each tumor immune cell clusters from **(E)** by individual replicates hashtagged from scRNA-seq. **(H)** CD8 T cells per milligram of tumor (left) or percent of CD44^+^PD1^+^ CD8 T cells from the draining lymph node (right) of mice bearing Res 499 tumors and pre-treated with or without anti- RANKL (n=7-8). Tumor and lymph nodes were harvested at day 15. P-values for survival are by log-rank test. For comparison between two groups, a two-sided T- test or Wilcoxon test is used for parametric or non-parametric data, respectively. See also Figure S2.

We next examined how anti-RANKL pre-treatment influences the overall immune cell composition of the tumor and draining lymph node (dLN). For this, we used single-cell RNA-seq (scRNA-seq) and hashtagging to assess the overall landscape from both tissue from multiple independent replicates of mice harboring Res 499 tumors and treated with dual ICB either with or without anti-RANKL pre-conditioning (Figure 2E-F and S2B-C). In the tumor, the only appreciable differences with anti-RANKL are an increase in CD8 T cells and a modest decrease in the DC3 subset of dendritic cells (Figure 2E and 2G). No significant changes in myeloid or neutrophil populations are observed in the setting of anti-RANKL pre-conditioning, in contrast to what is observed when anti-RANKL is given concurrent with ICB or when RANK is deleted from tumor cells (Ahern et al., 2017; Gómez-Aleza et al., 2020). In the dLN, a similar trend showing an increase in non-naïve CD8 T cells and cycling T cells expressing *Mki67* is evident after anti-RANKL (Figure 2F and S2D). Like the tumor, most other immune cells in the dLN are not significantly influenced by anti-RANKL, except for a decrease in activated B cells. Using flow cytometry, we confirmed that addition of anti-RANKL increases the proportion of CD8 T cells in tumors and the proportion of non-naïve CD8 T cells in the dLN, as measured by CD44^+^ PD1^+^ CD8 T cells (Figure 2H). Thus, RANKL blockade prior to tumor challenge and ICB does not lead to broad changes in the immune landscape but rather predominantly impacts properties of CD8 T cells relevant for ICB response.

### RANKL blockade mobilizes CD8 T cells against tumor self-antigens that are required for response

Immunotherapy against melanoma can result in on-target side-effects such as vitiligo due to an immune response against self-antigens shared by melanoma tumors and melanocytes (Byrne, 2011). Although only occurring in about 3% of melanoma patients, vitiligo is associated with a four-fold increase in survival (Teulings et al., 2015), suggesting the potential importance of tumor self-antigens in durable anti-tumor immune responses. Consistent with this notion, we observe that most mice harboring Res 499 melanoma that survive after sequential anti-RANKL plus anti-CTLA4 developed vitiligo between 32-40 days post-tumor challenge, an effect that was even more pronounced with anti-CTLA4 plus anti-PD1 (Figure 3A). In contrast, mice did not develop vitiligo when treated with ICB alone, treated with anti-RANKL without tumor challenge (Figure S3A), or treated with sequential anti-RANKL plus ICB and challenged with a Panc 4662 pancreatic tumor (Figure S3B). None of the anti-RANKL treated mice showed overt signs of additional autoimmunity but instead lived a natural lifespan despite treatment with ICB. Thus, the anti-tumor effect of RANKL blockade is associated with signs of enhanced autoimmunity against melanoma antigens after ICB.

**Figure 3.**
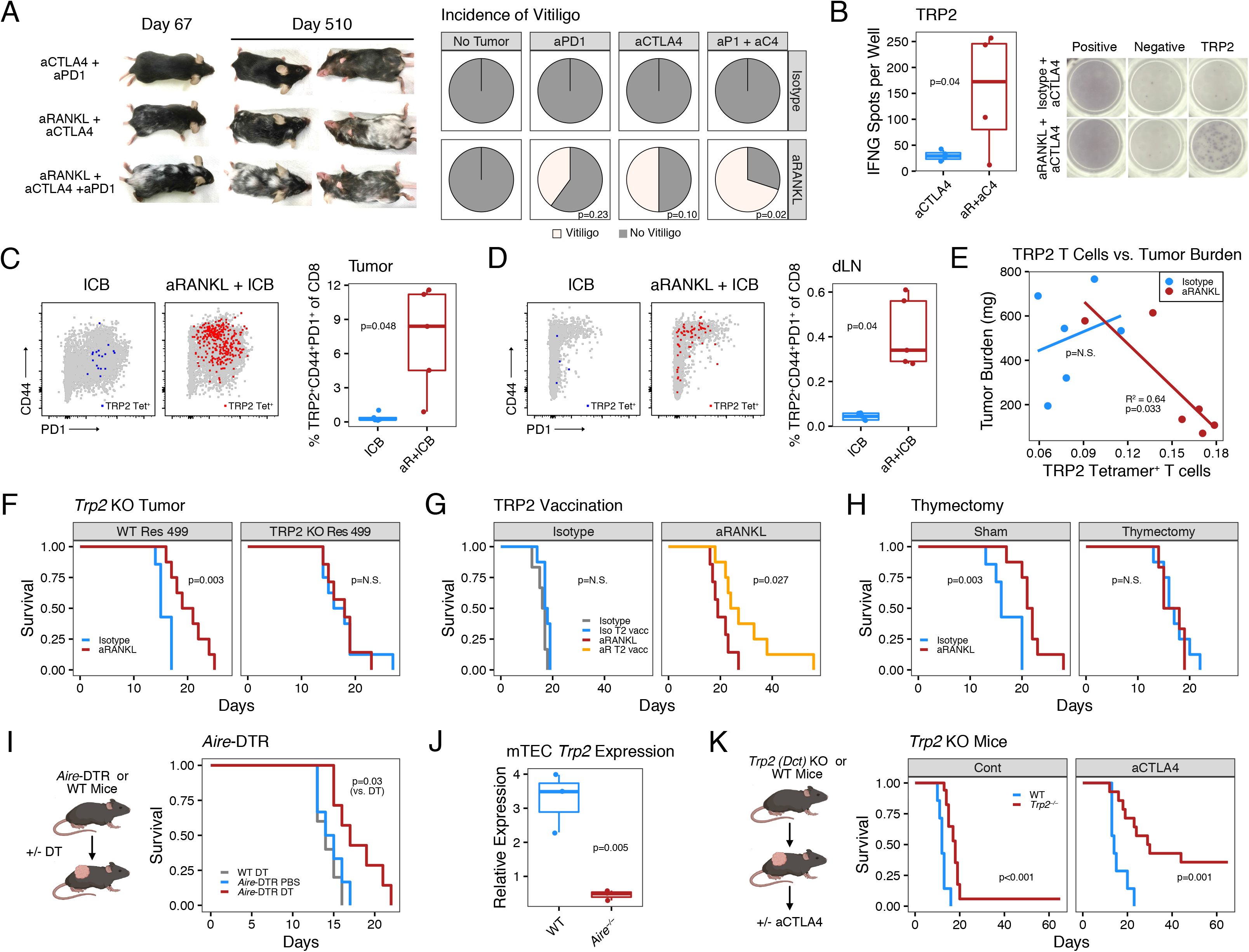
RANKL blockade promotes tumor-directed auto-reactivity through disruption of central tolerance. **(A)** Vitiligo in mice bearing Res 499 tumors after the indicated treatment. Shown are representative mice at days 67 and 510 and a pie chart of vitiligo incidence at day 60 (off-white). **(B)** Representative IFNG ELISpot image (right) and quantitation (left) of T cells reactive to TRP2_180-188_ peptide, a self-antigen epitope expressed by Res 499 melanoma. **(C-D)** Representative flow cytometry plots (left) and quantitation (right) of TRP2 tetramer^+^ CD8 T cells in the tumor **(C)** and draining lymph node **(D)** from mice treated with anti-CTLA4 + anti- PDL1 (ICB) with or without anti-RANKL pre-treatment. Data are day 15 post-tumor challenge. **(E)** Association between day 15 tumor burden and the frequency of TRP2 tetramer^+^ CD8 T cells in the draining lymph nodes in mice after pre-treatment with anti-RANKL or isotype control. Shown are regression lines through data from each treatment group. **(F)** Survival of mice bearing Res 499 tumors with or without TRP2 knockout after pre-treatment with anti-RANKL (n = 7-8). **(G)** Survival of mice bearing Res 499 tumors after pre-treatment with anti-RANKL and therapeutic vaccination with TRP2_180-188_ peptide (n=7-8). **(H)** Survival of thymectomized or sham-thymectomized mice pre-treated with anti-RANKL and challenged with Res 499 tumors (n=6-8). **(I)** Survival of control or *Aire*-diphtheria toxin receptor (DTR) mice bearing Res 499 tumors after treatment with diptheria toxin (DT) or PBS (n=5-7). **(J)** Relative expression of *Trp2* (*Dct*) in medullary epithelial cells (mTECs) from wild-type (WT) and *Aire* knock-out mice. **(K)** Survival of WT or *Trp2 (Dct)* KO mice bearing Res 499 tumors treated with or without anti- CTLA-4 (n=7-17). P-values for survival are by log-rank test. For comparison between two groups, a two-sided (or one-sided if changes in only one direction are expected) T-test or Wilcoxon test is used for parametric or non-parametric data, respectively. Linear regression is used to assess goodness of fit. See also Figure S3.

Given the high incidence of vitiligo in anti-RANKL treated mice, we screened known and predicted murine melanoma epitopes for reactivity using an ELISpot assay (Castle et al., 2012; Colella et al., 2000; Guevara-Patino et al., 2006; Overwijk et al., 1998; Wang et al., 1996). Splenocytes from mice challenged with Res 499 melanoma and treated with anti-RANKL plus anti-CTLA4 show reactivity to a TRP2_180-188_ peptide (Figure 3B), a previously characterized MHC-I-specific epitope derived from tyrosinase-related protein 2 (Bloom et al., 1997; Wang *et al*., 1996). To validate this, we stained paired splenocytes from our ELISpot assay with an H-2K^b^ tetramer loaded with TRP2_180-188_. This demonstrated an enrichment of a TRP2 tetramer^+^ population of CD8 T cells that are PD1^+^ Ki67^+^ GzmB^+^, consistent with the ELISpot screen (Figure S3C). Staining for TRP2 tetramer^+^ CD8 T cells in Res 499 tumors (Figure 3C) and the dLN (Figure 3D) revealed that these T cells are expanded in both tissues when anti-RANKL is added to ICB, suggesting specific activation and expansion upon tumor challenge. Thus, CD8 T cells against the tumor self-antigen TRP2 are systemically mobilized after RANKL blockade.

To elucidate whether the emergence of TRP2-reactive CD8 T cells after anti-RANKL contributes to the anti-tumor immune response, we examined the relationship between tumor burden and the percentage of TRP2 tetramer^+^ CD8 T cells from tumor dLNs. This revealed a significant negative correlation in response to anti-RANKL, whereas no correlation is observed in mice treated with an isotype control antibody (Figure 3E). Next, we directly tested the ability of TRP2 to serve as a dominant tumor rejection antigen by using CRISPR-Cas9 to knockout TRP2 expression in Res 499 melanoma (Figure S3D). Indeed, mice pre-conditioned with RANKL blockade no longer display delayed tumor outgrowth compared to isotype-treated counterparts (Figure 3F). Conversely, to examine if anti-RANKL enhances the generation of an anti-tumor responses against TRP2, we vaccinated Res 499 tumor-bearing mice with the TRP2_180-188_ peptide. Mice pre-conditioned with anti-RANKL exhibit prolonged survival after vaccination compared to isotype control (Figure 3G). Together, these data indicate that RANKL blockade promotes the utilization of TRP2 as a dominant tumor self-antigen for poorly immunogenic melanoma tumors.

### RANKL blockade interrupts central tolerance against self-antigens to improve immune checkpoint blockade

Based on the requirement for TRP2, we surmised that the primary mechanism of action of anti-RANKL is through the temporary depletion of *Aire*-expressing mTECs and interruption of central tolerance, resulting in the release of TRP2-reactive T cells that are normally deleted (Khan *et al*., 2014). However, RANKL and its receptor RANK are broadly expressed in other cell types and play important roles in bone metabolism, development, and immune regulation (Dougall et al., 1999; Green et al., 2002; Josien et al., 2000; Kong et al., 1999; Schramek et al., 2010). Therefore, we sought to confirm the extent to which effects of RANKL blockade on tumor response are due to interruption of central tolerance against TRP2 using a series of loss-of-function and genetic approaches. First, we examined the requirement for thymic T cell development by thymectomizing mice prior to RANKL blockade. Indeed, removal of the thymus ablated the effect of anti-RANKL compared to sham thymectomy, suggesting that new thymic emigrants are required for tumor response to RANKL blockade (Figure 3H). Second, to test if temporary depletion of *Aire*-expressing thymic mTECs analogous to our pre-conditioning with RANKL blocking antibody was sufficient to delay tumor outgrowth, we used *Aire*-diphtheria toxin receptor (DTR) mice that delete these cells upon administration of diphtheria toxin (Metzger *et al*., 2013). This revealed that mice injected with diphtheria toxin phenocopy those treated with anti-RANKL (Figure 3I). Next, we examined if *Aire* drives TRP2 expression in mTECs. Comparison of RNA sequencing data from wildtype to *Aire^-/-^* mice (St-Pierre et al., 2015) revealed that TRP2 expression in mTECs is regulated by *Aire* (Figure 3J). Lastly, to confirm the importance of this TRP2 expression and the role of negative selection in deleting high-affinity TRP2-reactive T cells, we implanted Res 499 tumors into *Trp2* (*Dct*) knockout mice and treated with ICB. As expected, loss of self-tolerance to TRP2 phenocopies the effects of anti-RANKL with and without ICB (Figure 3K). Together, these data suggest that the ability of anti-RANKL to enhance anti-tumor immunity is dependent on its role in disrupting central tolerance to TRP2 by depleting *Aire*-expressing mTECs.

Besides CD8 T cells, regulatory T cells (Tregs) are also selected based on their self-reactivity in the thymus, which is at least partly mediated by *Aire* (Jordan et al., 2001; Legoux et al., 2015; Metzger *et al*., 2013). To determine if diminished production or function of Tregs contribute to the delay in tumor outgrowth after anti-RANKL, we treated DEREG mice with diphtheria toxin to deplete Tregs after pre-conditioning with anti-RANKL. Treg depletion had a synergistic effect with RANKL blockade, indicating that Tregs are intact after anti-RANKL (Figure S3E). Moreover, we did not detect any differences in the percentages of Tregs in tumors and dLNs after anti-RANKL compared to control (Figure S3F). Thus, these results further corroborate that the dominant effect of anti-RANKL pre-conditioning prior to tumor challenge is the release of tumor-reactive CD8 T cells that are normally deleted by central tolerance.

### Interruption of central tolerance by RANKL blockade generates a tumor-reactive T cell repertoire with self-reactive features

In order to investigate how interfering with central tolerance using anti-RANKL alters the peripheral CD8 T cell repertoire against TRP2, we performed bulk TCR beta chain (TCRB) sequencing on peripheral TRP2 tetramer^+^ CD8 T cells from mice harboring Res 499 tumors and treated with ICB with or without anti-RANKL pre-conditioning (Figure S4A). First, the degree of similarity between the TRP2 repertoires of individual mice was assessed using a Jaccard index. This demonstrated generally higher similarity coefficients between mice treated with ICB alone compared to mice treated with anti-RANKL followed by ICB, indicating a less restricted TCR repertoire among mice treated with anti-RANKL (Figure 4A). Moreover, anti-RANKL treated mice have a more skewed repertoire as indicated by a higher Gini index, likely reflecting more pronounced proliferation among the top clonotypes (Figure 4B). Examination of TCRB V-gene usage reveals some repertoire biases after RANKL blockade (Figure S4B), but we observed no difference in the distribution of TCRB CDR3 lengths (Figure S4C). Together, these data suggest that RANKL blockade diversifies the TRP2 repertoire, generates unique TCR clonotypes, and enhances expansion of T cell clones.

**Figure 4.**
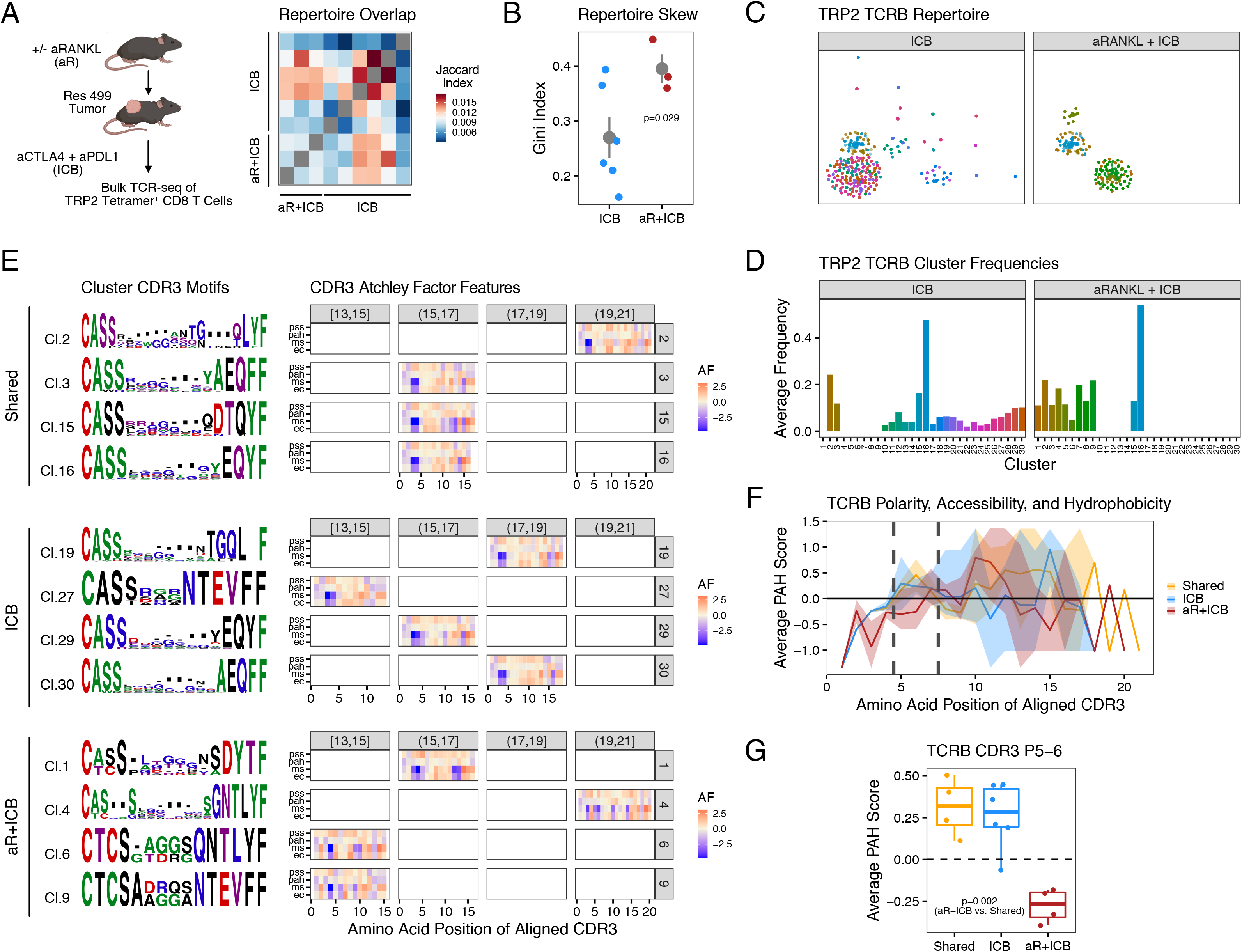
RANKL blockade alters the T cell repertoire and generates TCRs with self- reactive features. TCR beta chain (TCRB) sequencing was performed on TRP2 tetramer^+^ CD8 T cells from dLN and spleen of mice bearing Res 499 tumors and treated with anti-CTLA-4 plus anti-PDL1 (ICB) with or without anti-RANKL (aR) pre-treatment (n=3-6). **(A-B)** Shown is a heat map of pairwise Jaccard indices of TCRB CDR3 sequences to assess repertoire overlap **(A)** and Gini index to measure repertoire skewing **(B)**. **(C-D)** DeepTCR was used to characterize the TCRB CDR3 using supervised analysis to discern the TRP2 TCR repertoire from mice following the indicated treatment. Results were clustered and projected in low-dimensional UMAP space **(C)**. Average frequencies of the clusters for each treatment group are also shown **(D)** with the color matching the clusters shown in the UMAP. **(E)** Aligned TCRB CDR3 sequences from the most prevalent clusters found in the ICB-treated group, aRANKL + ICB group, or shared between both groups. Results for each cluster are shown using a sequence logo (row-wise). The average Atchley factor (AF) values that measure electrostatic charge (EC), molecular size (MS), polarity, accessibility, and hydrophobicity (PAH), and propensity for secondary structure (PSS) for each amino acid position in the aligned sequence are shown in the heat maps, which are additionally stratified by the indicated length of the aligned CDR3 (column-wise). **(F-G)** The average PAH scores for the aligned CDR3 from **(E)** is separately shown for each treatment group whereby the line indicates the average and the ribbon shows the range of PAH values for each amino acid position **(F)**. Values for positions 5 and 6 (P5-6), which are predicted peptide/MHC-I contact residues, were additionally averaged. Shown are results for shared clusters, clusters from ICB-treated mice, or clusters from mice treated with aRANKL + ICB **(G)**. For comparison between two groups, a two-sided T-test or Wilcoxon test is used for parametric or non-parametric data, respectively. See also Figure S4.

To gain deeper quantitative and qualitative insight into how anti-RANKL alters the TCR repertoire against TRP2, we used DeepTCR (Sidhom et al., 2021), a neural network deep learning method, to compare the expanded repertoire after ICB to the repertoire after anti-RANKL followed by ICB. For this, TCRB CDR3 sequences as well as VJ-gene usage was used for featurization followed by Monte Carlo simulation to develop the model (AUC = 0.76). The repertoire features learned by DeepTCR were then clustered and visualized by low-dimensional data projection using UMAP (Figure 4C). Among 30 total clusters are highly expanded clusters (e.g., cluster 16) that are shared between mice from both treatment groups (Figure 4D). In contrast, there are also multiple less prevalent clusters that are restricted to only one treatment group (Figures 4C-D). Therefore, to elucidate how group-specific clusters differ, top predictive TCRB CDR3 sequences from group-specific and shared clusters were aligned (see Methods). Then, each amino acid from this alignment was transformed into four Atchley factors that quantitates 1) electrostatic charge (EC), 2) molecular size (MS), 3) polarity, accessibility, and hydrophobicity (PAH), and 4) propensity for secondary structure (PSS). Examination of sequence logos reveal prominent differences in the CDR3s from anti-RANKL clusters, such as less restriction in the first four amino acid residues (Figure 4E, left). This is quantitatively demonstrated through the average Atchley factor values for the amino acids from the CDR3s in each cluster, grouped by CDR3 alignment length (Figure 4E, right). Also notable is a preference in the anti-RANKL plus ICB group for hydrophobic residues (leucine or valine) rather than negatively charged residues like glutamine two amino acids before the C-terminal end (e.g. LYF motif). This results in higher values for MS and EC but lower values for PAH near the C-terminus. Most prominent though, is an increase in average hydrophobicity (decreased PAH values) at amino acid positions 5 and 6 in the TCRB CDR3s from all anti-RANKL clusters compared to shared clusters or clusters from ICB-only (Figure 4F-G). Position 5 and 6 are thought to represent peptide-MHC contact residues, and hydrophobic amino acids at or near these positions can enhance TCR affinity to peptide-MHC and promote development of self-reactive T cells (Stadinski et al., 2016). Thus, RANKL blockade generates peripheral TRP2-reactive T cells with distinct TCR features that include specific changes in peptide-MHC contact residues linked to self-reactivity.

### T cells with self-reactive features generated by RANKL blockade escape exhaustion

Recent studies demonstrate that CD8 tumor-infiltrating lymphocytes (TILs) against tumor self-antigens in melanoma patients treated with ICB are typically exhausted and express low-affinity TCRs (Oliveira *et al*., 2021). Therefore, we sought to determine whether the altered T cell repertoire spawned by RANKL blockade might provide high-affinity TRP2-reactive CD8 TILs that can avoid exhaustion, potentially explaining how anti-RANKL improves ICB response. For this, we re-analyzed our scRNA-seq data along with accompanying single-cell TCR sequencing (scTCR-seq) data, focusing on CD8 T cells from Res 499 tumors and paired dLNs after ICB treatment. Examination of the transcriptional features of the TILs revealed six distinct clusters representing different CD8 T cell subsets (Figure 5A). To annotate these clusters, we used a mouse CD8 T cell exhaustion gene set and applied Gene Set Variation Analysis (GSVA) to first divide clusters into exhausted and non-exhausted states (Figure S5A). Next, for non-exhausted clusters, we repeated GSVA using gene sets for naïve-like (T_N_), memory-precursor (T_MP_), effector-memory (T_EM_), and terminal effector (T_TE_) CD8 T cell subsets. Annotations were additionally confirmed using well-known individual marker genes (Figure S5B). As expected, ICB monotherapy increases the proportion of CD8 T cells most strongly enriched for exhaustion genes (T_EX.2_) (Figure 5B). In contrast, anti-RANKL pre-conditioning prior to ICB decreased the frequency of this population, while increasing the proportion of T_MP_ or T_EM_ cells. The identity of this T_EM_ cluster is corroborated by expression of T_EM_-associated transcription factors such as *Prdm1*, *Id2*, and *Stat4* (Figure 5C). Anti-RANKL also increases the frequency of CD8 TILs that expresses effector-memory and terminal effector genes, annotated as T_EM_TE_ (Figure 5B-C). Similar analysis of non-naïve CD8 T cells in the corresponding dLNs (Figure 5D) revealed that the most pronounced change after anti-RANKL is an increase in T cells belonging to the two **<u>S</u>**cRNA-seq lymph **<u>N</u>**ode clusters SN2 and SN6 (Figure 5E) that are also enriched in the memory-precursor gene and effector-memory genes sets (MP and EM), and corresponding genes such as *Il7r, Id2* and *Stat4* (Figure 5F). Thus, these data suggest that RANKL blockade promotes tumor-infiltration of CD8 T cells that avoid T cell exhaustion and develop memory-precursor and effector-memory features.

**Figure 5.**
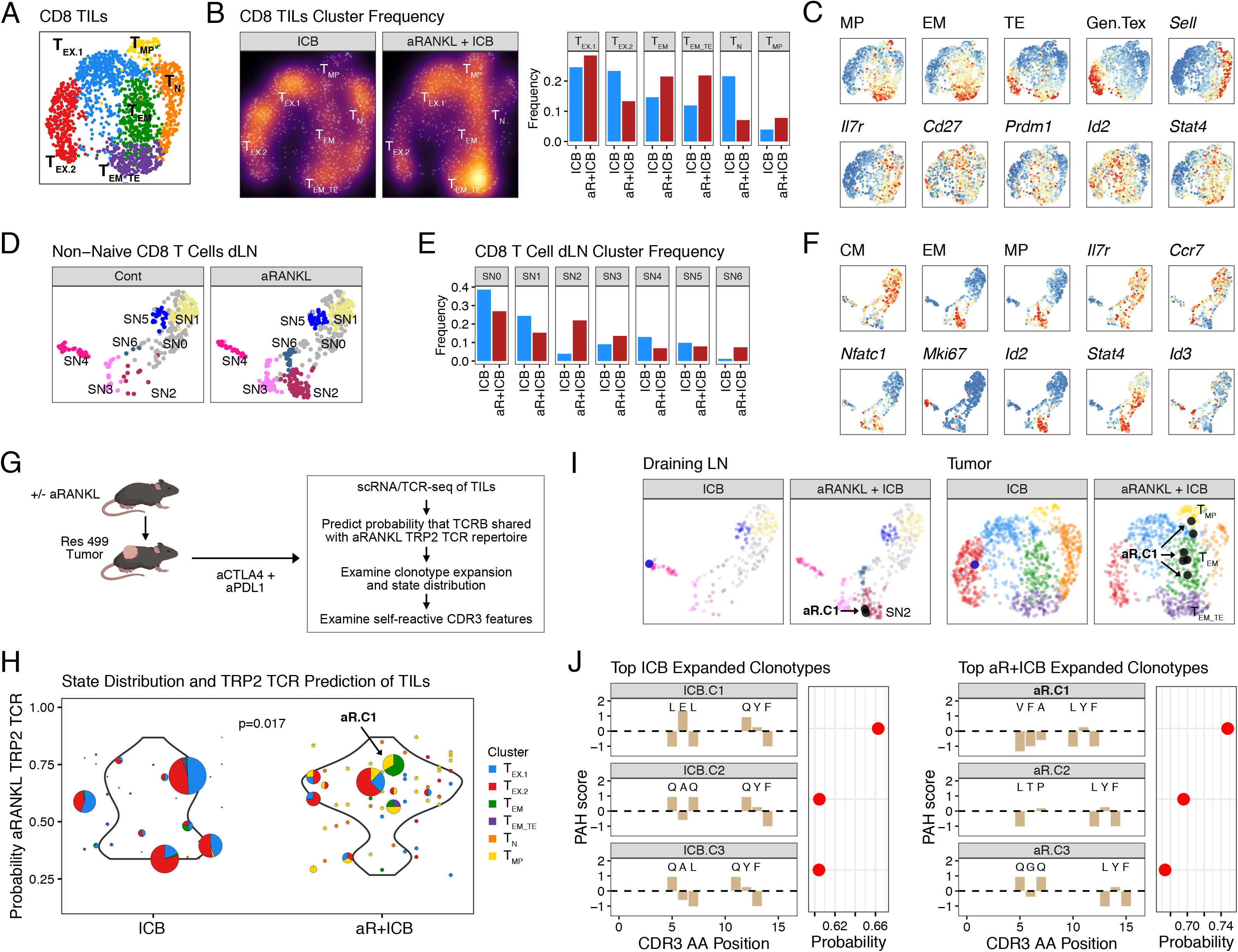
T cells generated by RANKL blockade avoid exhaustion and give rise to memory-precursor and effector-memory populations in response to ICB. Single-cell RNA-seq was performed from Res 499 tumors and draining lymph nodes (dLN) of mice treated with anti-CTLA4 + anti-PDL1 (ICB) with or without anti-RANKL pre-treatment at day 15 post-tumor challenge (n=4, pooled). **(A-C)** UMAP plot of the CD8 T cells from Res 499 tumors. Cluster annotations are shown **(A)** along with density overlay and plot of relative frequencies for each cluster **(B)**. Relative expression of the indicated markers are overlaid on the UMAP plot **(C)**. **(D-F)** UMAP plot of CD8 T cells from the dLN. Cluster annotations are shown **(D)** along with plot of relative frequencies for each cluster **(E)**. Relative expression of the indicated markers are overlaid on the UMAP plot **(F)**. **(G-I)** Single-cell TCR-seq of the TCRB and concurrent scRNA-seq was performed on TILs from (A). The probability that the TCRB is shared with the TRP2 CD8 T cell repertoire generated after aRANKL + ICB (aRANKL TRP2 TCR) was predicted using a Monte Carlo DeepTCR model (AUC=0.76), and other properties of the clonotype were examined **(G)**. Shown is the aRANKL TRP2 TCR probability along with the CD8 T cell subtype distribution for expanded clonotypes from the indicated groups **(H)**. Clonotypes that are shared between paired dLN and tumor for each treatment group are overlaid on the UMAP plots **(I)**. Highlighted is clonotype aR.C1 that belongs to dLN cluster SN2 and T_MP_ and T_EM_ clusters in the tumor. **(J)** PAH scores of key amino acids of the TCRB CDR3 from expanded TIL clonotypes from (H). The three expanded clonotypes with a TCR with the highest probability of being shared with the TRP2 CD8 T cell repertoire generated after ICB alone (left) or after aRANKL + ICB (right) are shown. Bar plots show PAH scores for amino acid positions 5-7 and the last three C-terminal amino acids in the aligned CDR3. The abutting dot plot on the right-margin indicates the TRP2 TCR repertoire probability. Highlighted in bold is clonotype aR.C1 from the aRANKL + ICB group to facilitate cross-reference to (H) and (I). A KS test is used determine differences in DeepTCR prediction result. See also Figure S5.

We next analyzed the accompanying scTCR-seq data to evaluate the effect of RANKL blockade on individual CD8 T cell clonotypes. Minimal apparent clonal expansion, as determined by the proportion of unique TCRB CDR3 sequences, occurs in the dLNs of mice treated with ICB, whereas 15% of clonotypes in mice treated with anti-RANKL plus ICB are non-unique and hence expanded (Figure S5C). Furthermore, almost all these expanded dLN clones belong to cluster SN2 that is enriched in MP and EM genes (Figure S5D). Compared to the dLN, clonality assessment of TCRB sequences from TILs revealed greater expansion in tumors from mice treated with ICB compared to ICB with anti-RANKL (62% and 43%) (Figure S5E). However, without anti-RANKL, these expanded TIL clonotypes almost exclusively reside in T_EX.1_ and T_EX.2_ exhausted clusters, while expanded clonotypes from anti-RANKL treated mice show a much larger proportion in the T_MP_ cluster (Figure S5F). This expansion of memory-precursor clonotypes is accompanied by accumulation of T_EM_ and T_EM_TE_ subtypes, possibly due to subsequent differentiation. Thus, anti-RANKL promotes priming and clonal expansion of T cells with effector-memory rather than exhausted features.

Next, we sought to assess whether the clonotypes that give rise to T_EM_ or T_EM_TE_ in the tumor share key features with the TRP2 T cell repertoire generated by RANKL blockade, and whether these T cells can be traced to T_MP_ clonotypes and priming events in the dLN. For this, we used our DeepTCR model trained to distinguish features of the anti-RANKL TRP2 T cell repertoire to predict the likelihood that TIL clonotypes belong to this repertoire (Figure 5G). Indeed, most expanded clonotypes from anti-RANKL treated tumors are predicted to arise from the anti-RANKL TRP2 repertoire, as indicated by higher and skewed probability scores compared to scores from CD8 T cells treated with ICB alone (Figure 5H). Analysis of how each clonotype distributes among CD8 T cell subsets indicates that clonotypes predicted to belong to the anti-RANKL repertoire (probability score > 50%) are predominantly T_MP_ or expanded from T_MP_. Notably, among extensively expanded clonotypes (Figure 5H, larger circles), the clonotype with the highest probability score after anti-RANKL plus ICB, denoted clonotype aR.C1, is comprised of both T_MP_ and T_EM_ subsets, suggesting a lineage relationship. Moreover, clonotype aR.C1 also belongs to the expanded MP/EM-like dLN cluster SN2, demonstrating lymph node priming likely preceded tumor infiltration (Figure 5I). Analysis of the TCRB CDR3 from clonotype aR.C1 confirmed that it contains hydrophobic amino acids (low PAH score) at positions 5 through 7 and the LYF triplet at the C-terminal end (Figure 5J, right plots), both of which characterize the TRP2 T cell repertoire after RANKL blockade (Figure 4E-G). In contrast, these self-reactive features are less prominent in expanded clonotypes with lower probabilities. Neither of these CDR3 features are seen in the top expanded and primarily exhausted CD8 TIL clonotypes predicted to resemble TRP2-specific T cells generated by ICB alone (Figure 5J, left plots). Together, these data suggest that preconditioning with anti-RANKL prior to ICB treatment promotes dLN proliferation of CD8 T cells with T_MP_ features that can then infiltrate and expand into dominant TILs that avoid exhaustion. These T cells are predicted to be TRP2-reactive with TCR features indicative of self-reactivity.

### Effector-memory T cells generated by RANKL blockade exhibit increased TCR affinity for tumor self-antigens

To more directly demonstrate that effector-memory dLN and tumor CD8 T cells generated by RANKL blockade possess TCRs with high-affinity against TRP2, we sought to develop a flow cytometry antibody panel that can discriminate T cell subsets and approximate TRP2 TCR affinity by tetramer MFI (Crawford et al., 1998; Yee *et al*., 1999). Examination of individual gene markers from the scRNA-seq data revealed that intermediate to high expression of *Pdcd1*, *Tbx21, Gzmb, Cd44*, *Itgal*, and *Cxcr3* and low to intermediate expression of *Sell* approximate T cells with MP/EM-like features (Figure S6A). Using the dLN and tumors from mice implanted with Res 499 melanoma cells and treated with either ICB or sequential anti-RANKL plus ICB, we then used antibodies against proteins encoded by these markers for flow cytometry. Dimensionality reduction and clustering of expression intensities revealed five distinct **<u>F</u>**low cytometry-derived lymph **<u>N</u>**ode (FN) clusters (Figures 6A-B and Figure S6B) with varying intensities for the TRP2 tetramer (Figure 6C). Although expectedly rare, T cells with high TRP2 tetramer intensities are more abundant across independent replicates from the anti-RANKL plus ICB compared to ICB-only group (Figure 6D). Moreover, these T cells are primarily found in dLN cluster FN5 (Figure 6E, above grey dotted line) and resemble activated T cells expressing PD1, Ki67, and GzmB (compare Figure 6E right to Figure 6B). To confirm these TRP2 tetramer-high T cells also resemble cluster SN2 dLN T cells with EM/MP-like features, we sorted TRP2 tetramer^+^ CD8 T cells from the dLN and spleen of mice with Res 499 tumors treated with anti-RANKL plus ICB. Single-cells were then plated and individual TRP2 tetramer intensity values were captured prior to scRNA-seq. This revealed that compared to TRP2 tetramer MFI-low cells, MFI-high cells have greater enrichment of genes differentially expressed by the MP/EM-like dLN cluster SN2 (Figure 6F). In contrast, differentially expressed genes from other clusters such as SN0 show a decrease in MFI-high cells, while genes from other dLN clusters are not significant.

**Figure 6.**
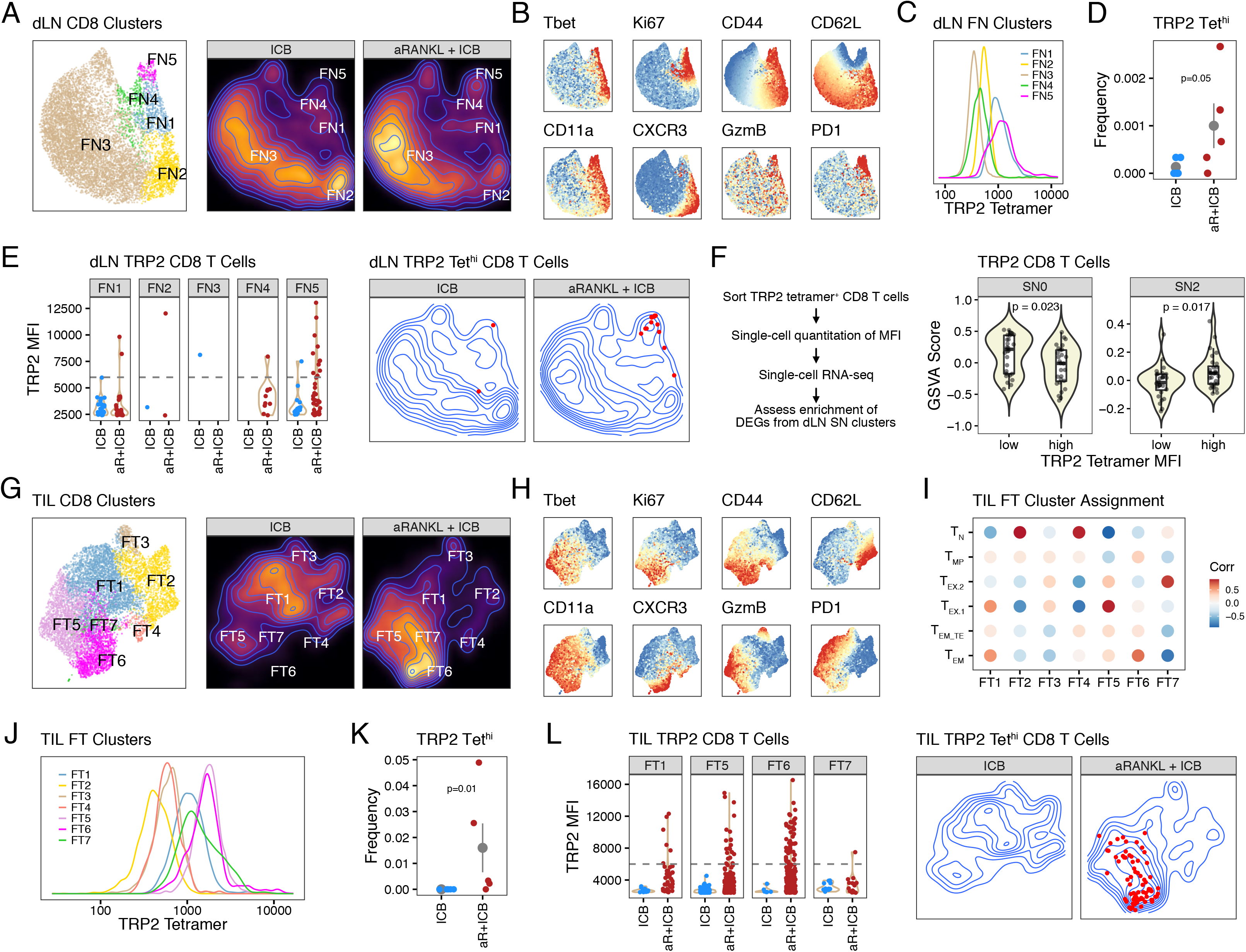
Memory-precursor and effector-memory T cells generated after RANKL blockade exhibit high-affinity for the tumor self-antigen TRP2. Flow cytometry for markers selected to approximate scRNA-seq clusters from the tumor and dLN was performed on mice bearing from Res 499 tumors and treated with anti-CTLA4 + anti- PDL1 (ICB) with or without anti-RANKL pre-treatment at day 15 post-tumor challenge. **(A-B)** UMAP plot of the CD8 T cells from the dLN (n=5, pooled). Cluster annotations are shown **(A)** along with overlay of relative fluorescent intensities of the indicated markers **(B)**. **(C-E)** TRP2 tetramer fluorescent intensities for each dLN FN cluster **(C)** along with the relative frequency of T cells that are TRP2 tetramer MFI-high (Tet^hi^) for each replicate from the indicated treatment group **(D)**. The distribution of TRP2 tetramer-positive CD8 T cells across the dLN clusters **(E)** with the TRP2 Tet^hi^ population indicated above the grey dotted line (left). Also shown (right) are these TRP2 Tet^hi^ cells overlaid on the UMAP from (A). **(F)** TRP2 tetramer-positive CD8 T cells from dLN and spleen of mice bearing Res 499 tumors and treated with aRANKL + ICB were sorted and plated as single-cells. The tetramer MFI for individual cells was determined by fluorescence microscopy prior to scRNA-seq. Single-cell enrichment for genes differentially expressed across the dLN SN clusters from Figure 5D was determined. Shown are the enrichment scores for the genes from the indicated SN clusters for TRP2 tetramer MFI-low versus MFI-high CD8 T cells. **(G-H)** UMAP plot of the CD8 T cells from the tumor (n=5, pooled). Cluster annotations are shown **(G)** along with overlay of relative fluorescent intensities of the indicated markers **(H)**. **(I)** Pairwise correlations between average FT cluster expression of markers shown in **(H)** with corresponding RNA expression from scRNA-seq clusters shown in Figure 5A. **(J-L)** TRP2 tetramer fluorescent intensities for each CD8 TIL FT cluster **(J)** along with the relative frequency of T cells that are TRP2 tetramer MFI-high (Tet^hi^) for each replicate from the indicated treatment group **(K)**. The distribution of TRP2 tetramer-positive CD8 T cells across the TIL clusters **(L)** with the TRP2 Tet^hi^ population indicated above the grey dotted line (left). Also shown (right) are these TRP2 Tet^hi^ cells overlaid on the UMAP from (G). For comparison between two groups, a two-sided (or one-sided if changes in only one direction are expected) T-test or Wilcoxon test is used for parametric or non-parametric data, respectively. See also Figure S6.

Flow cytometry analysis of tumor-infiltrating CD8 T cells demonstrated seven **<u>F</u>**low cytometry-derived **<u>T</u>**umor (FT) clusters (Figures 6G-H). Among these clusters, cluster FT6 is more prominent after anti-RANKL plus ICB versus ICB alone (Figures S6C). Correlating average per cluster expression of the surface markers to expression of their corresponding genes from the scRNA-seq-defined clusters demonstrated that cluster FT6 most resembles T_EM_ cells, and to a lesser degree T_MP_ (Figure 6I). Cluster FT6 TILs also exhibit high TRP2 tetramer intensity (Figure 6J). Indeed, across independent replicates CD8 T cells with high TRP2 tetramer intensity are more frequent after addition of anti-RANKL (Figure 6K) and predominantly reside in the EM/MP-like cluster FT6 (Figure 6L, right contour plot, and above grey dotted line in left violin plot). In contrast, with ICB alone TRP2 tetramer MFI-high TILs are nearly absent.

Altogether, these results suggest that RANKL blockade generates TRP2-reactive CD8 T cells with TCRs that not only have self-reactive features but also are high-affinity. These CD8 T cells are primed in the dLN and preferentially give rise to TILs belonging to T_MP_ and T_EM_ subsets, linking TCR affinity and lymph node priming to the circumvention of T cell exhaustion.

### Memory-precursor CD8 T cells that acquire TCRs with self-reactive features after RANKL blockade are enriched for NFAT/AP-1 genes

TCR activation that results in cooperation between NFAT and AP-1 transcription factors promotes the differentiation of effector-memory rather than exhausted T cells (Martinez et al., 2015). Therefore, we sought to assess whether acquisition of high-affinity TRP2 TCRs by MP/EM-like CD8 T cells is linked to greater TCR signaling and NFAT/AP-1 gene expression after tumor infiltration. Utilizing Nur77-GFP reporter mice, which assesses TCR signaling strength (Ashouri and Weiss, 2017; Moran et al., 2011), greater Nur77 activity is observed in CD8 T cells from Res 499 tumors after anti-RANKL plus ICB versus ICB alone (Figure 7A), which is consistent with differential gene expression and gene set enrichment analysis as well (Figure S7A-B). In particular, TRP2 tetramer^+^ CD8 T cells exhibit significantly higher Nur77 activity in the tumor after RANKL blockade (Figure 7B), suggesting high-affinity TRP2-reactive TCRs are accompanied by greater TCR signaling. Based on these observations, we next utilized scRNA/TCR-seq to examine Nur77 gene expression of TILs and relate this with features of TRP2 TCRs generated by anti-RANKL (Figure 7C). Indeed, among T_MP_ and T_EM_ cells there is a significant correlation between *Nur77* (*Nr4a1)* expression and the probability that that T cells possess a TCR similar to TRP2 T cells resulting from RANKL blockade (Figure 7D). When this probability in T_MP_ cells exceeds 50%, increasing *Nur77* is also accompanied by enrichment of NFAT/AP-1 genes (Figure 7E). Importantly, the positive correlation between NFAT/AP-1 gene enrichment and TCR features is particularly strong in T_MP_ TILs after RANKL blockade (Figure 7F, slope of regression lines; Figure S7C), suggesting that the ability of anti-RANKL to augment NFAT/AP-1 genes depends on TILs acquiring anti-RANKL-associated TCR features. Together, these data corroborate that RANKL blockade generates high-affinity and self-reactive T cells against TRP2 that undergo greater TCR activation in the tumor after ICB. These high-affinity cells exhibit coordinated expression of transcription factors that may help early precursor T cells to circumvent exhaustion, enabling the development of effector-memory TILs.

**Figure 7.**
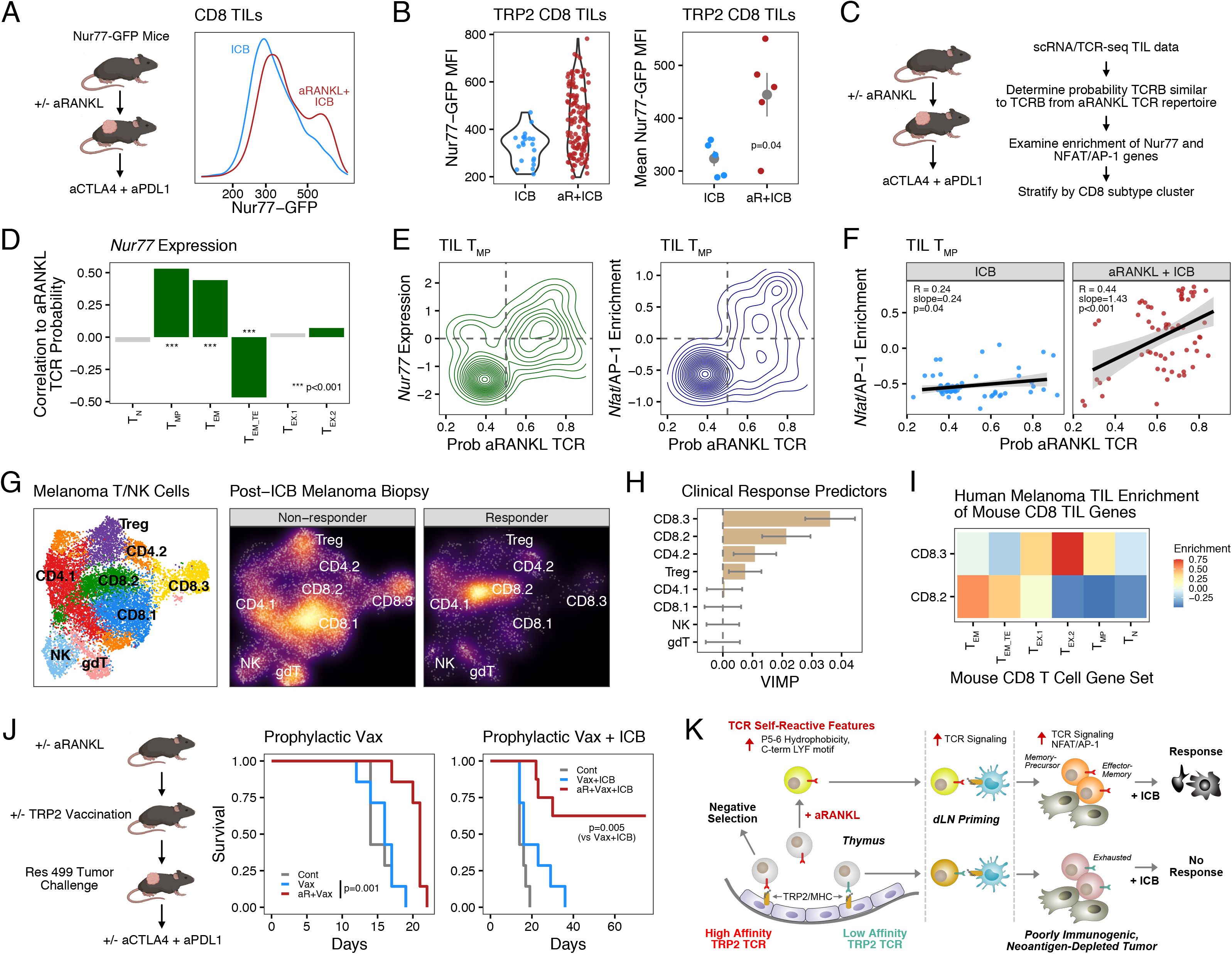
High-affinity TRP2 CD8 T cells generated by RANKL blockade display strong TCR signaling, enrichment for NFAT/AP-1 genes, and transcriptional features that predict clinical ICB response. **(A-B)** TCR signaling strength measured using Nur77-GFP mice bearing Res 499 tumors after treatment with anti-CTLA4 + anti-PDL1 with or without anti-RANKL pre-treatment. Shown are the fluorescent intensities for the Nur77-GFP from all CD8 TILs **(A)** and the distribution from TRP2 tetramer-positive TILs **(B)** along with the mean Nur77-GFP MFI for TRP2 tetramer- positive TILs for each replicate. **(C-F)** Single-cell TCR-seq of the TCRB and concurrent scRNA-seq was performed on TILs from day 15 Res 499 tumors of mice treated with anti-CTLA4 + anti-PDL1 (ICB) with or without anti-RANKL pre-treatment (n=4, pooled). The probability that the TCRB is shared with the TRP2 CD8 T cell repertoire generated after aRANKL + ICB (aRANKL TRP2 TCR) was predicted using a DeepTCR model, and other properties of the clonotype were examined **(C)**. Shown is the correlation between the aRANKL TRP2 TCR probability and *Nur77* (*Nr4a1*) expression for each of the indicated CD8 T cell subtypes **(D)**. For T_MP_ TILs, which have the highest correlation, the association between the aRANKL TRP2 TCR probability **(E)** and *Nur77* expression (left) or NFAT/AP-1 gene enrichment (right) is separately plotted along with NFAT/AP-1 enrichment stratified by treatment group **(F)**. **(G)** T and NK cells from human melanoma biopsies from patients treated with ICB (anti-PD1 +/- anti-CTLA4) were clustered and annotated (left). The density of the cluster frequencies in non-responders and responders are shown overlaid on a UMAP plot (right) (n=48, pooled by response category). **(H)** The relative frequencies for the T/NK cell clusters were used to predict response using random forest (error rate 0.42). The top predictors were determined by variable importance score (VIMP). Shown are VIMP averages and standard deviation from Monte Carlo simulation. **(I)** The human melanoma T/NK clusters from (G) were examined for enrichment of genes differentially expressed by the mouse Res 499 melanoma TIL clusters from Figure 5A. Shown are GSVA enrichment scores for the two CD8 T cell clusters with the top VIMPs for clinical response prediction. **(J)** Survival of mice receiving prophylactic vaccination with TRP2_180-188_ peptide after pre- conditioning with anti-RANKL (left) or with subsequent ICB-treatment (right) after challenge with Res 499 tumors. **(K)** Summary model of the mechanism of action for RANKL blockade. P-values for survival are by log-rank test. For comparison between two groups, a two-sided T-test or Wilcoxon test is used for parametric or non-parametric data, respectively. Linear regression is used to examine goodness of fit. See also Figure S7.

### RANKL blockade improves efficacy of prophylactic cancer vaccination and promotes T cell transcriptional features that predict clinical ICB response

Since most tumor-reactive T cells in human melanoma have been reported to be exhausted (Oliveira *et al*., 2021), we sought to determine whether TILs that transcriptionally resemble the TILs generated after RANKL blockade are associated with improved ICB response compared to exhausted TILs. For this, we created gene sets from differentially expressed genes that distinguish each CD8 T cell subset in Res 499 tumors after ICB with or without anti-RANKL. Then, we used random forest supervised learning on scRNA-seq data from 48 human melanoma patients treated with ICB (Sade-Feldman et al., 2018) to identify subsets of TILs in post-treatment tumors that are associated with response or non-response. This demonstrated that the top two melanoma TIL clusters that predict ICB outcome are both CD8 T cell clusters, with cluster CD8.3 primarily found in non-responders and cluster CD8.2 in responders (Figure 7G-H). While cluster CD8.3 is most enriched in genes differentially expressed by the T_EX.2_ exhausted subset from Res 499 tumors, cluster CD8.2 is most enriched in genes from T_EM_, the subset promoted when anti-RANKL is combined with ICB (Figure 7I). Thus, RANKL blockade generates CD8 TILs with transcriptional features that are associated with ICB response in mice and humans.

Besides improving tumor response after ICB, using anti-RANKL to transiently provide the T cell repertoire with high-affinity T cells against tumor self-antigen could potentiate prophylactic cancer vaccination. To test this, we vaccinated mice against TRP2_180-188_ during the window of mTEC depletion after anti-RANKL but delayed tumor challenge until *Aire* expression is restored (Khan *et al*., 2014). Consistent with our therapeutic vaccination results (Figure 3G), prophylactic vaccination with TRP2 peptide prolonged survival in anti-RANKL treated but not isotype-treated mice (Figure 7J). Similarly, prophylactic vaccination after anti-RANKL, but not prophylactic vaccination alone, markedly improves subsequent ICB therapy against Res 499 tumors (Figure 7J, right). Collectively, these data suggest that generation of high-affinity CD8 T cells against self-antigen using RANKL blockade can improve prophylactic cancer vaccination.

## DISCUSSION

Most common human cancers may be insufficiently antigenic to mount an effective tumor-directed T cell response after ICB treatment. Cancer immunoediting that rids tumors of clones expressing strong neoantigens and central tolerance that prunes the T cell repertoire of high-affinity T cells against potential tumor self-antigens are likely two major factors that contribute to poor tumor immunogenicity. In this study, we show that RANKL blockade can overcome such limitations to ICB efficacy by temporarily interrupting central tolerance and spawning high-affinity T cells against self-antigens (Figure 7K). Some of these T cells that are normally deleted possess TCRs with distinct autoreactive features capable of targeting self-antigens expressed by the tumor, such as TRP2. Upon addition of ICB, these T cells proliferate in the lymph node, exhibit high levels of TCR signaling, and support a transcriptional program in memory-precursors that is linked to the circumvention of T cell exhaustion. These T_MP_ cells go on in the tumor to adapt an effector-memory phenotype that helps mount a self-antigen-directed response against neoantigen-depleted tumors.

Our study provides insight into the role of TCR affinity, T cell exhaustion, and efficacy of ICB. Immune checkpoint blockade therapies can only galvanize T cells that are available in the repertoire, which are generally depleted of high-affinity self-reactive T cells due to central tolerance (Hogquist and Jameson, 2014). Recent studies that measure both TCR affinity (or avidity) and transcriptional readouts of T cell fate and function reveal that the majority of exhausted TILs from human tumors are tumor-reactive, while effector-memory TILs have TCRs that recognize non-tumor antigens (Oliveira *et al*., 2021). Exhaustion of tumor-reactive TILs occurs regardless of whether the T cells recognize neoantigens or tumor self-antigens. Interestingly, although T cells against neoantigens were high-avidity and T cells against tumor self-antigens were low-avidity, neoantigens were found to be expressed at low levels on cancer cells while tumor self-antigens were expressed at high levels. Such findings suggest that a mismatched relationship between TCR avidity and antigen availability might result in blunted TCR signaling that favors the development of exhaustion. This inverse relationship is likely a consequence of cancer immunoediting and host central tolerance, raising the question of whether increasing TCR affinity can avoid T cell dysfunction and improve ICB response. Indeed, high-affinity TCRs have been linked to successful adoptive cell therapies and ICB in lung cancer (Caushi et al., 2021; Sim et al., 2020). In our study, we disrupt the apparent inverse relationship between TCR affinity and antigen levels that is imposed by central tolerance through use of RANKL blockade *in vivo* without *a priori* knowledge of the target antigen or antigens. This strategy may be particularly useful for patients with lesser-known HLA haplotypes or tumors for which appropriate target antigens have not been characterized. Our results suggest that therapeutically and acutely generating T cells with high-affinity against tumor self-antigens *in vivo* can circumvent T cell dysfunction. Whether other factors besides intrinsic properties of the TCR also contribute to the prevention of T cell exhaustion requires additional investigation.

Our data suggests that natural repertoires, absent of transgenic or adoptively transferred cells, are poor at rejecting solid tumors without the addition of ICB. These include repertoires resulting from RANKL blockade, direct depletion of *Aire*-expressing mTECs, and those for which TRP2 is not a self-antigen. This is likely influenced by various immunosuppressive mechanisms typical of solid tumors and may also be due in part to the rapid and aggressive outgrowth of our melanoma tumor model. However, it also suggests that peripheral mechanisms of tolerance are efficient at suppressing high-affinity autoreactive responses. Similarly, we do not observe acute “off-target” autoimmunity or exacerbation of known irAEs with RANKL blockade despite widespread incidence of “on-target” autoimmunity in the form of vitiligo. Indeed, exactly how *Aire* deficiency manifests itself with regard to immune responses and autoimmunity outside of autoimmune polyendocrinophathy-candidiasis-ectodermal dystrophy syndrome (APECED) has proven to be nuanced (Davis, 2015; 2016; Mathis and Benoist, 2009). The limited toxicity we observe may be partially explained by tissue-seeding of long-lived perinatal Tregs with enhanced suppressive capacity that may guard against autoimmunity (Guerau-de-Arellano et al., 2009; Yang et al., 2015). After disrupting central tolerance through RANKL blockade, a precipitating or threshold priming event such as tumor challenge or vaccination may be required in tandem with ICB treatment to drive a corresponding auto-response.

One possible limitation of the use of RANKL blockade as therapeutic strategy to improve tumor immunogenicity is the ability of cancer patients to produce new T cells. Thymic involution decreases T cell output with age (Goronzy et al., 2015); however, measurable and significant thymic output can continue into older adulthood (Thome et al., 2014; Vrisekoop et al., 2008; Wallace et al., 2004). Furthermore, many cancer types such as breast cancer, melanoma, and pediatric tumors can develop at younger ages before complete thymic involution either stochastically or due to genetic predisposition (Cruz-Rodriguez et al., 2016; Krepischi et al., 2012; Navid et al., 2016). In fact, there is clinical evidence that cancer patients might benefit from RANKL blockade from reports of patients treated with denosumab, a humanized RANKL blocking antibody originally approved for the treatment of osteoporosis. For example, a recent prospective clinical trial of premenopausal women with early breast cancer found that pre- operative treatment with denosumab is associated with a significant increase in CD8 T cell infiltration into tumors despite having no effect on the proliferation or survival of tumor cells (Gómez-Aleza et al., 2020). Interestingly, case reports and retrospective studies suggest potential synergy between denosumab and ICB therapy with no apparent increase in irAEs (Afzal and Shirai, 2018; Angela et al., 2019; Liede et al., 2018; Smyth et al., 2016). Notably, RANKL blockade may have other effects that contribute to suppression of tumor growth (Ahern et al., 2017; Gómez-Aleza et al., 2020; Ming et al., 2020). These complementary mechanisms of action likely do not depend on thymic output of T cells.

RANKL blockade may improve other treatment modalities besides ICB that would benefit from the provision of high-affinity T cells against tumor self-antigen. For instance, tumor self-antigens have been desirable vaccine targets due to near-ubiquitous expression on cancer cells. For melanoma, tumor self-antigens are expressed in 93-95% of tumors irrespective of stage (Barrow et al., 2006) with higher expression correlated with increased ICB response in patients with neoantigen-poor tumors (Lo et al., 2021). However, such cancer vaccinations against self-antigens have shown limited efficacy either alone or in combination with ICB (Hodi *et al*., 2010; Rosenberg et al., 2004). Presumably, either therapeutic or prophylactic vaccination against self-antigens is also limited by central tolerance, both of which can also be overcome through RANKL blockade. Prophylactic vaccination may be particularly attractive to those with known familial risk for developing cancer and can be given before concerns about thymic involution occur.

In summary, our studies provide key insight into the interplay between central and peripheral mechanisms of tolerance in anti-tumor immunity and delineate the mechanism by which RANKL blockade sensitizes refractory tumors to ICB therapy. Moreover, our work elucidates a clinically available, synergistic, out-of-the-box therapy potentially capable of altering the T cell repertoire to favor durable anti-tumor responses that may enable otherwise unresponsive patients to benefit from ICB and other immunotherapies.

## ACKNOWLEDGMENTS

E.D. and A.J.M. were supported by the Melanoma Research Alliance. A.J.M. was supported by the Parker Institute for Cancer Immunotherapy, the Breast Cancer Research Foundation, the Mark Foundation for Cancer Research, and grants from the NIH (1P01CA210944-01) and Department of Defense (W81XWH-17-1-0264). E.D. was supported by a grant from the NIH (F31CA213915) and by the Patel Scholar Award for Cancer Research from the University of Pennsylvania School of Medicine. We thank the laboratory of Dr. Andrea Facciabene for assistance with ELISpot assays. We also thank the NIH Tetramer Core Facility (contract number 75N93020D00005) for providing H-2K^b^ mouse TRP2 and SIINFEKL tetramers. Some figure illustrations were created with BioRender.com.

## AUTHOR CONTRIBUTIONS

Conceptualization, E.D., M.S.A., and A.J.M.; Methodology, E.D., O.O., R.R., and A.J.M.; Formal Analysis, E.D., R.R., O.O., and A.J.M.; Investigation, E.D., O.O., and H.D.; Writing –Original Draft, E.D. and A.J.M.; Writing – Review & Editing, E.D., M.S.A., and A.J.M.; Funding Acquisition, E.D., M.S.A., and A.J.M.; Project Administration, E.D. and A.J.M.; Supervision, A.J.M.

## SUPPLEMENTAL FIGURES

**Figure S1.**
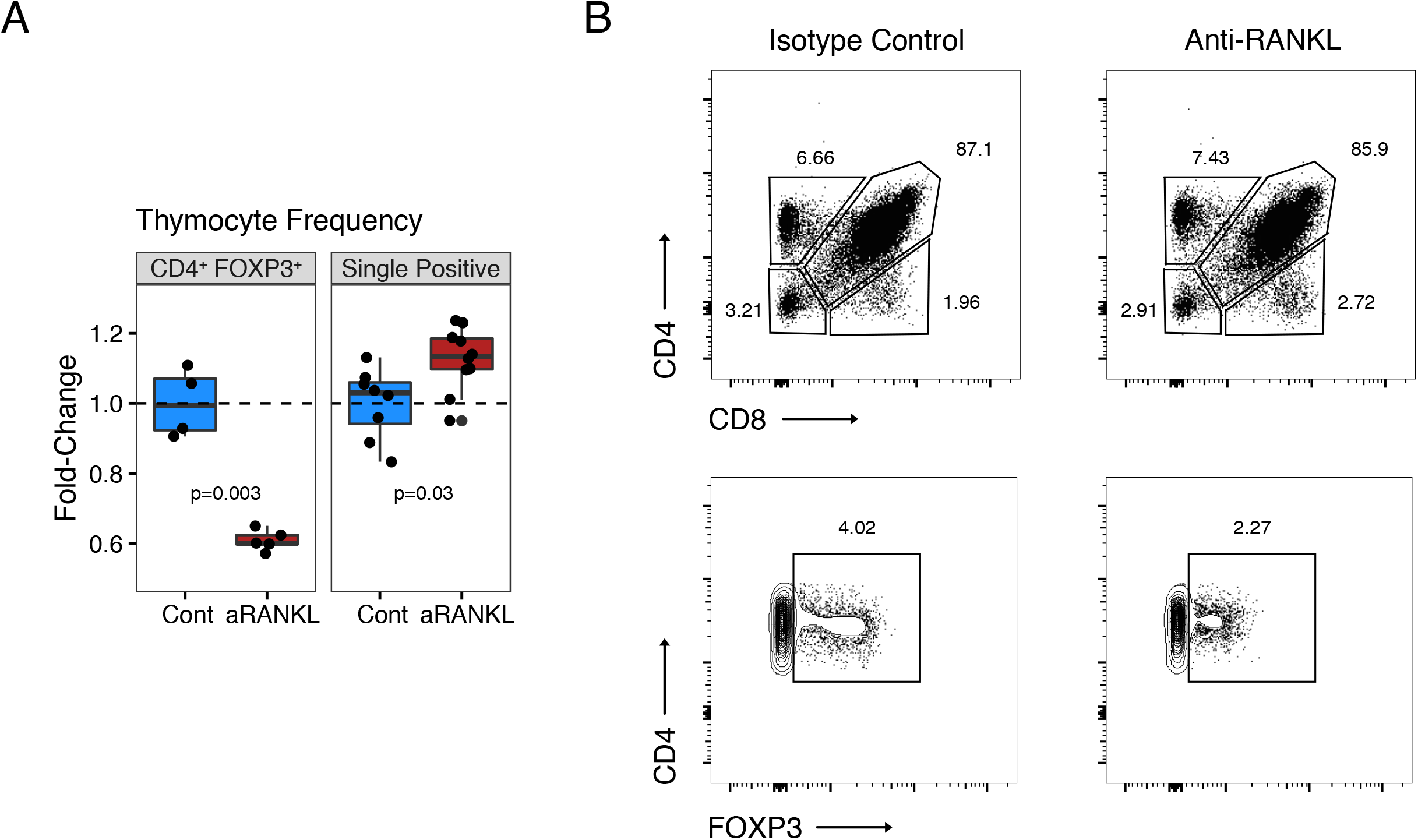
RANKL blockade temporarily alters T cell development in the thymus (Related to Figure 1) **(A)** Fold-change in indicated thymocyte populations from mice treated with anti-RANKL or isotype control. Values are normalized to average of isotype control. The single positive population includes both CD4 and CD8 cells. **(B)** Representative flow cytometry plots of thymocytes from (A). Percentage of cells in each of the indicated gates are shown.

**Figure S2.**
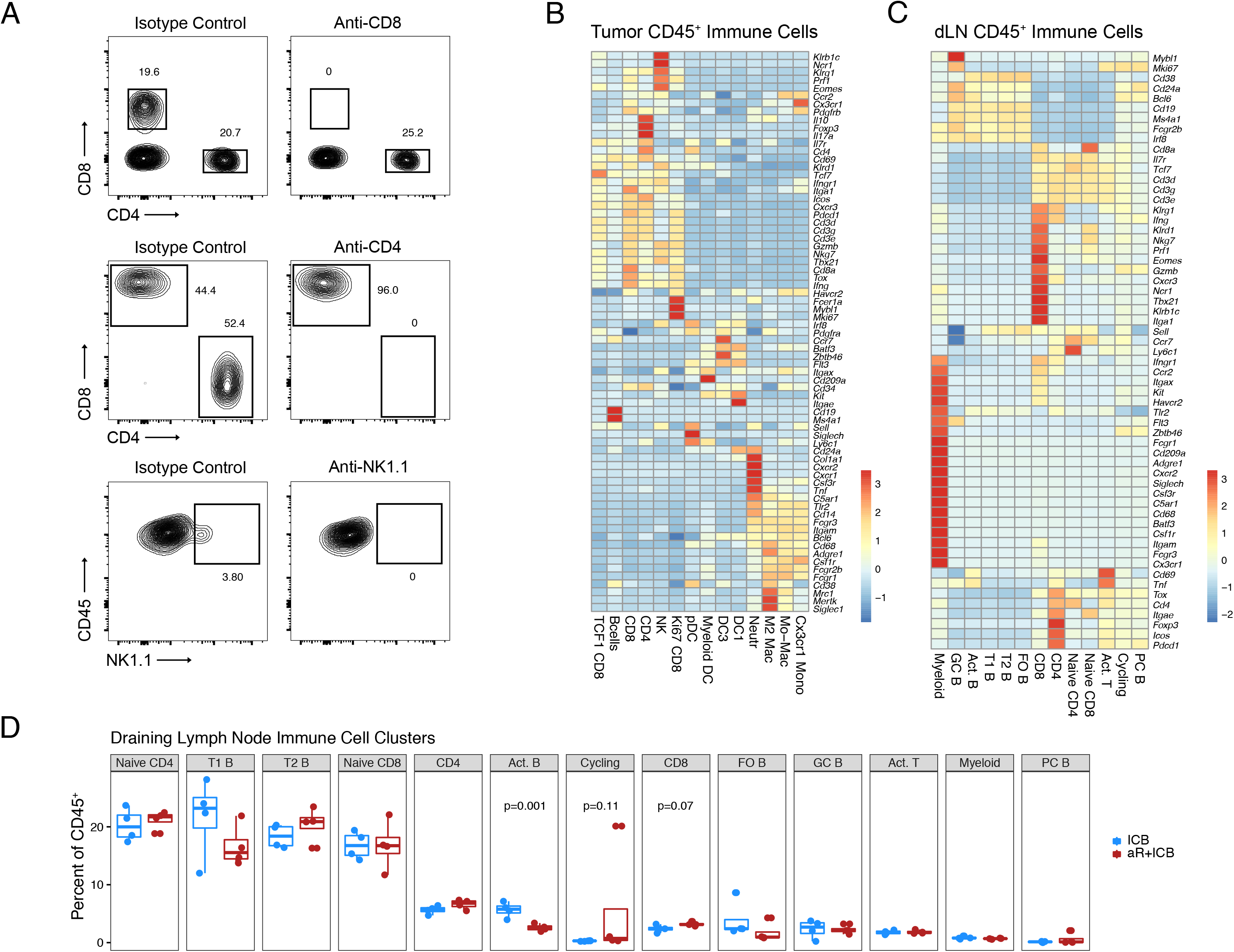
Immune population requirements and features associated with RANKL blockade (Related to Figure 2) **(A)** Representative flow cytometry plots of CD4 and CD8 T cells and NK1.1^+^ cells after treatment with a depleting antibody or isotype control. Percentage of cells in each of the indicated gates are shown. **(B-C)** Scaled expression of the indicated marker genes for CD45^+^ immune cells from the tumor **(B)** or draining lymph node **(C)**. **(D)** Percentage of CD45^+^ cells in each draining lymph node cluster shown in Figure 2F by individual replicates hashtagged from scRNA-seq.

**Figure S3.**
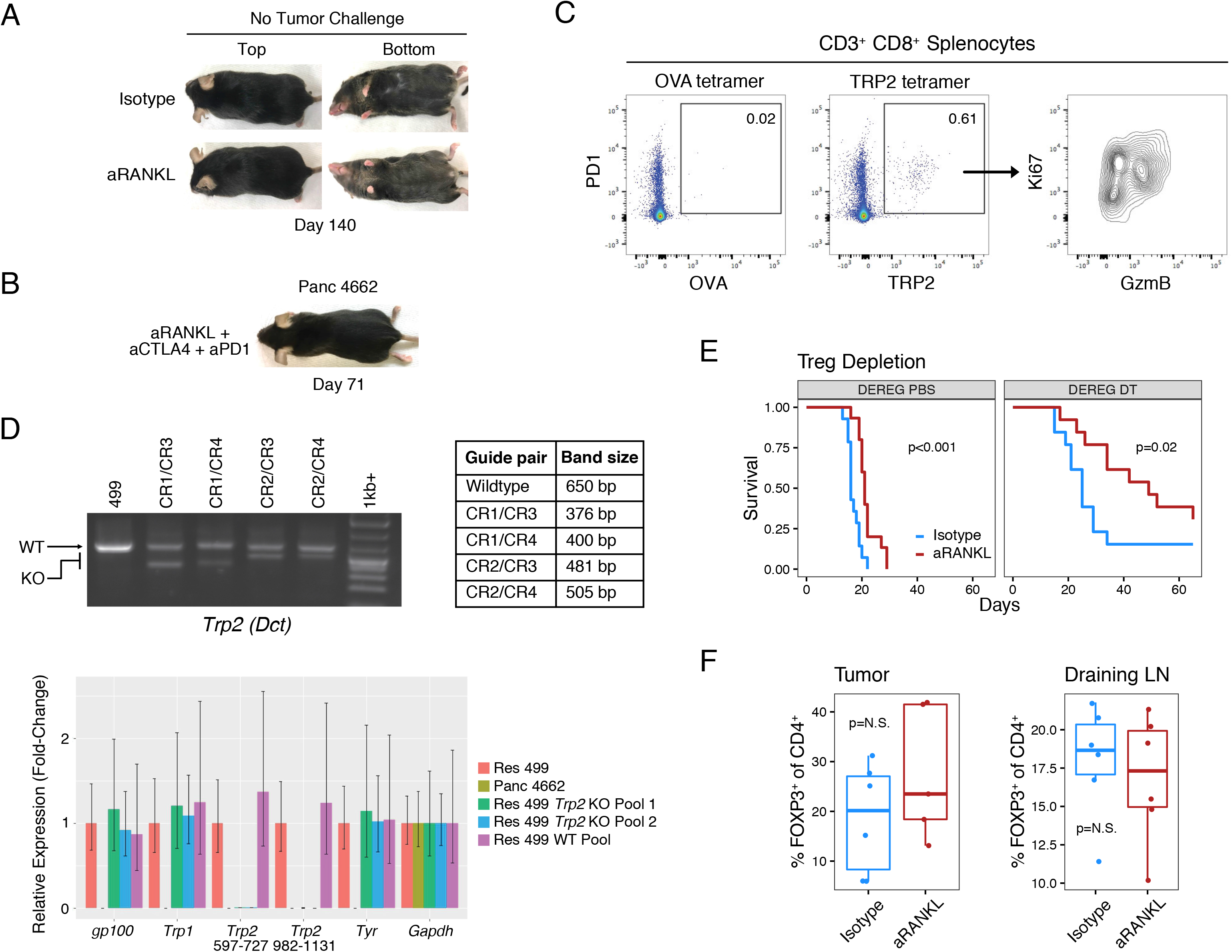
RANKL blockade promotes expansion of CD8 T cells against TRP2 (Related to Figure 3) **(A)** Vitiligo in representative C57BL/6 mice bearing Res 499 melanoma tumors treated with or without anti-RANKL at 140 days post-tumor challenge. **(B)** Representative mouse bearing a Panc 4662 pancreatic tumor at day 71 after treatment with anti-RANKL with anti-CTLA4 and anti-PD1. **(C)** Representative TRP2 tetramer-positive CD3^+^ CD8^+^ splenocytes from mice bearing Res 499 tumors and treated with anti-RANKL and anti-CTLA-4. TRP2 tetramer-positive cells were additionally assessed for Ki67 and GzmB (right plot). An OVA tetramer was used as a control. **(D)** PCR amplification products with primers specific to the *Trp2* (*Dct*) gene locus using DNA from sorted Res 499 cancer cells after CRISPR knockout of TRP2. Expected position of PCR products from wild type and knockout cells are shown next to the gel along with a table of anticipated PCR fragment sizes. Also shown is a bar plot of the relative expression by qRT-PCR of *Trp2* (using two separate primer sets) and other melanoma self-antigens from pooled Res 499 TRP2 WT and KO cells and a Panc 4662 pancreatic cell line used as a negative control. **(E)** Survival of *Foxp3*-DTR mice bearing Res 499 tumors and treated with diphtheria toxin (DT) or PBS control (n = 13-15). P-values for survival are by log-rank test. **(F)** Percent FOXP3^+^ CD4^+^ T cells in the tumors and draining lymph nodes of mice pre-treated with anti-RANKL prior to Res 499 ICB-resistant B16-F10 tumor challenge (n = 6).

**Figure S4.**
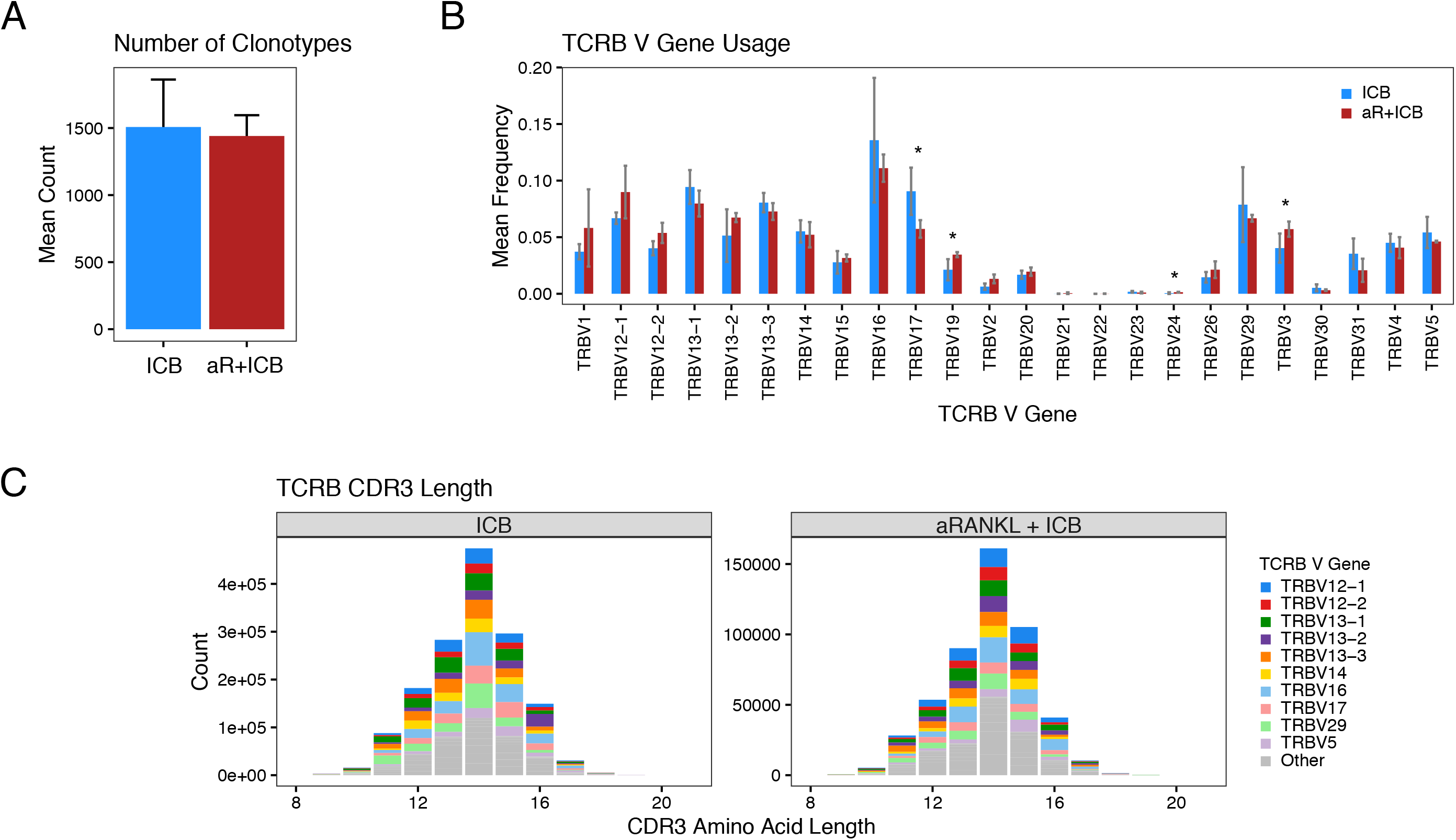
TCR features of the TRP2-reactive CD8 T cell repertoire after RANKL blockade (Related to Figure 4) **(A-C)** Average number of clonotypes **(A)**, average frequency of TCRB V genes **(B)**, and TCRB CDR3 amino acid lengths **(C)** from peripheral TRP2-tetramer positive CD8 T cells sorted from mice treated with anti-CTLA-4 plus anti-PDL1 (ICB) with or without anti-RANKL. Error bars in (A) are standard error of the mean and in (B) are standard deviations. For comparison between two groups, a two-sided T-test is used (*, p ≤ 0.05).

**Figure S5.**
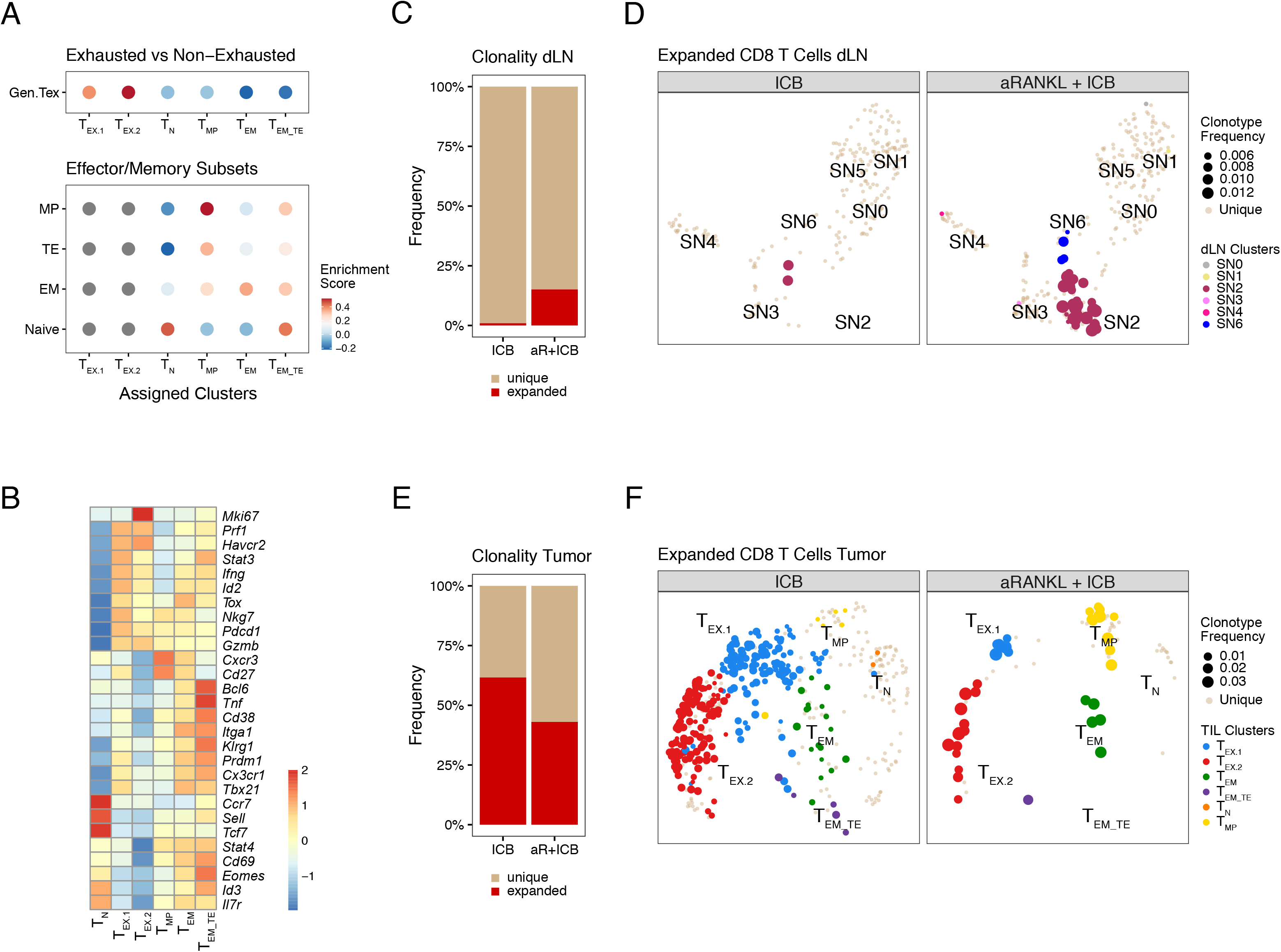
Expansion and features of CD8 T cells from draining lymph node and tumor after RANKL blockade (Related to Figure 5) **(A)** Enrichment of gene sets for CD8 T cell subsets (y-axis) for each annotated CD8 T cell cluster (x-axis) from tumors shown in Figure 5A. Top plot shows enrichment for a general T cell exhaustion gene set, while bottom plot shows enrichment for naïve-like, effector, or memory subset genes in the non-exhausted clusters. **(B)** Heat map of scaled expression values for the indicated marker genes across the annotated CD8 TIL clusters from (A). **(C-D)** Frequency of expanded clonotypes (non-unique) from tumor draining lymph nodes **(C)** and the T cells belonging to these expanded clonotypes projected on the UMAP from Figure 5D **(D)**. The clonotype frequency for these expanded T cells and the cluster membership is indicated on the UMAP. The non-expanded clonotypes (unique) are also shown on the UMAP. **(E-F)** Frequency of expanded clonotypes (non-unique) from the tumor **(E)** and the T cells belonging to these expanded clonotypes projected on the UMAP from Figure 5A **(F)**. The clonotype frequency for these expanded T cells and the cluster membership is indicated on the UMAP. The non-expanded clonotypes (unique) are also shown on the UMAP.

**Figure S6.**
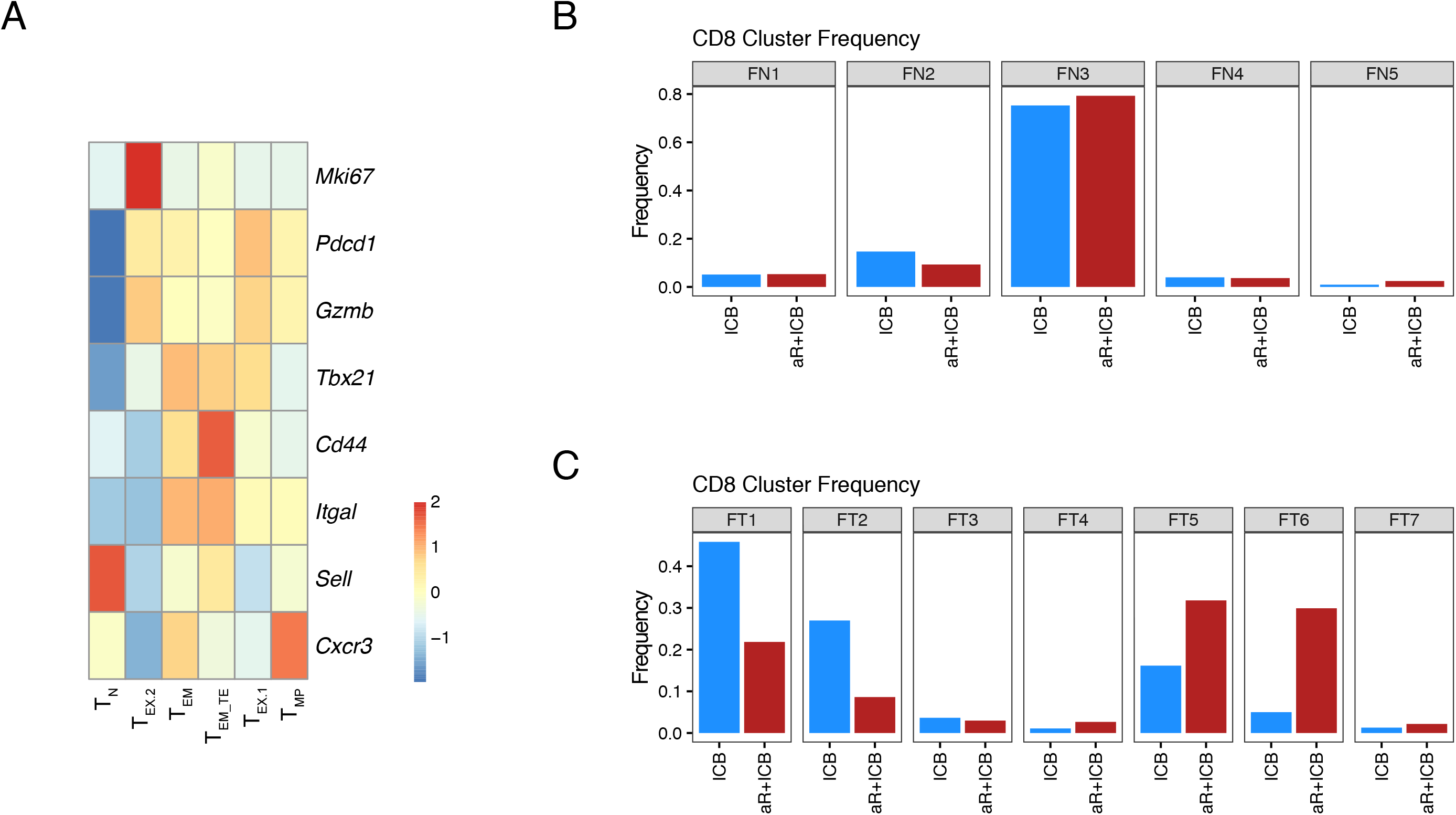
Frequency and features of CD8 T cells from draining lymph node and tumor after RANKL blockade (Related to Figure 6) **(A)** Heat map of the indicated marker genes across the annotated CD8 TIL clusters from Figure 5A used to inform markers used for flow cytometry. **(B-C)** Frequency of CD8 T cells in the draining lymph node **(B)** and tumor **(C)** from mice treated with ICB with or without anti-RANKL. Frequencies and T cell cluster membership was determined by flow cytometry using protein markers for the genes shown in (A).

**Figure S7.**
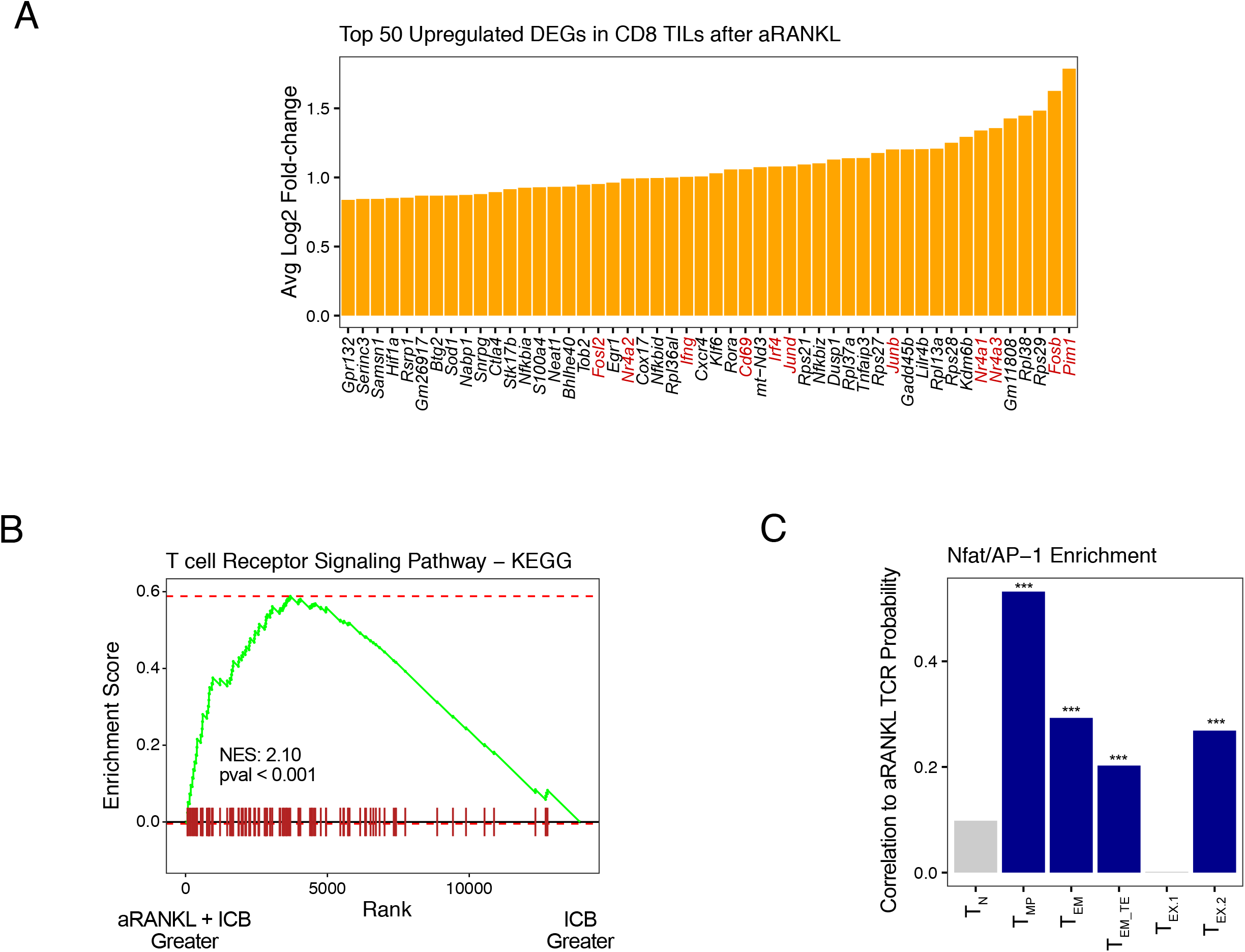
Gene expression changes in tumor-infiltrating CD8 T cells after RANKL blockade (Related to Figure 7) Single-cell TCR-seq of the TCRB and concurrent scRNA-seq was performed on TILs from day 15 Res 499 tumors of mice treated with anti-CTLA4 + anti-PDL1 (ICB) with or without anti-RANKL pre-treatment. **(A-B)** Top 50 differentially upregulated genes **(A)** and enrichment of genes associated with TCR signaling **(B)** in tumor-infiltrating CD8 T cells after anti-RANKL plus ICB versus ICB alone. Select genes associated with TCR signaling and activation are highlighted in red in (A). The normalized enrichment score (NES) and p-value for KEGG TCR signaling gene set enrichment are indicated in (B). **(C)** The probability that the CD8 T cell TCRB is shared with the TRP2 CD8 T cell repertoire generated after aRANKL + ICB was predicted using a DeepTCR model (Figure 5C). Shown is the correlation between this probability and NFAT/AP-1 gene enrichment for each CD8 T cell subset.

## MATERIALS AND METHODS

### Experimental Model and Subject Details

#### Mice

Five- to seven-week-old C57BL/6 (000664), BALB/c (000651), and DEREG (32050-JAX) mice were obtained from the Jackson Laboratory (Bar Harbor, ME). Six-week-old C57BL/6 mice were thymectomized or sham-thymectomized at the Jackson Laboratory. Aire-DTR mice (Metzger *et al*., 2013) were transferred from the University of California, San Francisco. *Dct* KO mice (Guyonneau et al., 2004) were rederived from sperm using female C57BL/6 mice from the Jackson Laboratory. Nur77-GFP reporter mice (Moran *et al*., 2011) were a gift from Michael Cancro. David Feldser and Paula Oliver generously provided Rag1 KO mice. Mice were maintained under pathogen-free conditions. All animal experiments were performed according to protocols approved by the Institutional Animal Care and Use Committee (IACUC) of the University of Pennsylvania.

#### Cell Lines

Res 499 resistant B16-F10 melanoma, Res 237 resistant TSA breast cancer were derived and cultured as previously described (Twyman-Saint Victor *et al*., 2015). The Panc 4662 pancreas cancer cell lines were derived as previously described (Bayne *et al*., 2012).

#### Peptides

OVA, mgp100, and TRP2 peptides were purchased from Anaspec, Inc. (Fremont, CA). Custom peptides were ordered from Eunoia Biotech (Wynnewood, PA) and ProImmune, Inc. (Sarasota, FL).

### Method Details

#### *In vivo* mouse studies

Mice were administered either anti-RANKL (IK22/5) or isotype control (2A3) antibody intraperitoneally at 100ug/injection three-times per week for two weeks. Tumor injection and treatment schedule were performed as previously described (Twyman-Saint Victor *et al*., 2015). Mice that had achieved complete responses were rechallenged on both flanks between days 60-90 days post-tumor challenge and monitored for relapse. *In vivo* blocking antibodies anti-CTLA4 (9H10), anti-PD1 (RMP1-14, J43), and anti-PDL1 (10F.9G2) were administered at 200ug/injection on days 5, 8, and 11. 200ug/dose of CD8 (53-6.7), CD4 (GK1.5), and NK1.1 (PK136) depleting antibodies were administered on days -2, 0, and every 3 days thereafter. Diphtheria Toxin (Sigma) was injected intraperitoneally at 25-50ng/g.

#### IFNγ ELISpot assay

Ninety-six well MAIP plates (Millipore) were coated with purified rat anti-mouse IFNγ (clone R4-6A2, BD Pharmingen). Splenocytes were harvested on day 15 post-tumor inoculation, red blood cells removed with ACK buffer, and plated at 5x10^5^ cells/well in duplicate. Plates were incubated overnight at 37°C with 1 mg/ml suspension of each peptide, washed with 0.05% Tween 20/PBS, incubated overnight at 4°C with biotin-conjugated rat anti-mouse IFNγ (clone XMG 1.2, Pharmingen), and developed with NBT/BCIP (Pierce). Spots were counted using an automated ELISpot reader.

#### Flow cytometry and cell sorting

Tumors, draining lymph nodes, and spleens were harvested on day 15 after tumor challenge. Single-cell suspensions were generated and treated with ACK Lysis Buffer (Life Technologies) to remove red blood cells as previously described (Twyman-Saint Victor *et al*., 2015). Cell viability was established by Live/Dead Fixable Aqua Dead Cell Stain Kit (Life Technologies) or DAPI staining (Thermo Fisher). TruStain fcX (Biolegend) was used to block non-specific antibody staining. Surface staining was performed for 30 min at 4 degrees. Tetramer (NIH Tetramer Facility) staining was performed at 37 degrees for 45 min. Foxp3 fixation/permeabilization kit (eBioscience) was used for intracellular staining. To determine immune infiltration, tumor sample weight was recorded prior to staining and the entire sample was collected by flow cytometer. Flow cytometry was performed on the LSR II (BD) and analyzed using *FlowJo* software (TreeStar), *FlowCore* (v1.52.1), and *FlowSOM* (v1.16.0). The FACSAria II (BD) was used for cell sorting.

#### Vaccination

Mice were injected subcutaneously with a total volume of 200ul containing 100ug TRP2_180-188_ peptide and 50ug poly(I:C) (InvivoGen) formulated in in PBS. Mice were vaccinated 4 days prior to tumor challenge and again 7 days later as previously described (Castle *et al*., 2012). For prophylactic vaccination, mice were injected twice, one week apart, 7 weeks prior to tumor challenge.

#### CRISPR gene targeting

Gene targeting by CRISPR/Cas9 was accomplished by dual- transfection of pX458 (GFP) and pX459 (puromycin) Cas9-expression vectors (Addgene, 48138; Addgene, 62988) containing paired RNA guides flanking the TRP2_180-188_ epitope. GFP^hi^ cells were collected and subsequently selected using puromycin to isolate co-transfected cells. Successful targeting of *Trp2* (*Dct*) was determined by two independent PCR-based screening methods in addition to qPCR analysis of single-cell clones. Knock-out cells or control cells were pooled separately in equal numbers to maintain the diversity of the parental population. The guide RNA and screening oligo sequences used are listed below:

- *Dct* gRNA #1: GGACCGGCCCCGACTGTAATCGG
- *Dct* gRNA #2: TGCGTGGTGATCACGTAGTCTGG
- *Dct* gRNA #3: GGCTTACCGCGCATCATTAGAGG
- *Dct* gRNA #4: TGATGCGCGGTAAGCCAATTTGG
- *Dct*-F: TTCTCGGAGGGTTCTCTGTG
- *Dct*-R: ACAGGCGGTGTTTGGTAATC
- *Dct*KO-intC1-F: GCCTTAGACCTGGCCAAGAAGA
- *Dct*-982F: TTCGAGACACATTATTAGGTCCAG
- *Dct*-1131R: GCAACGCAAAGGACTCATTG

#### Bulk and single-cell tetramer^+^ T cell sequencing

Spleens and draining lymph nodes were harvested and disrupted to form single-cell suspensions. CD8 T cells were enriched with the EasySep™ Mouse CD8 T Cell Isolation Kit (Stem Cell Technologies). Cells were stained, resuspended in DAPI at 0.25 μmol, and live tetramer^+^ T cells were collected using a FACSAria II (BD), excluding residual B220^+^, CD4^+^ and NK1.1^+^ cells. Bulk TCR sequencing was performed using the SMARTer® Mouse TCR a/b Profiling Kit (Takara Bio). Libraries were sequenced using the 600-cycle MiSeq Reagent Kit v3 (Illumina) with paired-end, 2 x 300 base pair reads and analyzed with *MiXCR* (https://github.com/milaboratory/mixcr). For single-cell analysis, cells were injected into the C1 Single-Cell Auto Prep IFC for mRNA Seq (5-10um) and run on the C1 platform with the SMART-Seq® v4 Ultra® Low Input RNA Kit (Fluidigm). Calcein AM (Thermo Fisher) was used to establish cell viability and tetramer-APC intensity was determined. Tandem gene expression and TCR sequence determination was performed using *TraCeR* (Stubbington et al., 2016).

#### Single-cell RNA, TCR, and feature sequencing

Tumors and draining lymph nodes were harvested on day 15 post-tumor challenge and single-cell suspensions were generated as described above. Total live CD45^+^ leukocytes were isolated on the FASCAria II (BD), excluding dead cells by DAPI staining. Single-cell libraries were prepared using the Single Cell Immune Profiling v1 with Feature Barcode technology protocol and corresponding reagents (10x Genomics). Libraries were sequenced using the NextSeq 500/550 v2.5 High Output Kit (Illumina). TotalSeq™-C anti-mouse Hashtag Antibodies 1-4 (BioLegend) were used to differentiate between tumor samples.

### Quantification and Statistical Analysis

#### Analysis of tumor growth, survival, and statistical analysis

Mice were randomly assigned into age and sex-matched treatment groups. Tumor volume was determined by caliper and calculated using the formula: *L* × *W* ^2^ × 0.52; in which *L* is defined as the longest dimension and *W* is its perpendicular. For survival studies, an event was defined as death, tumor ulceration, or when tumor burden reached a size of 1.5 cm in any dimension to minimize morbidity, in accordance with our study protocol. Differences in survival were determined for each group using the Kaplan-Meier method from *survival* (v3.1-8) and pairwise comparisons were calculated by logrank test. Separations in tumor growth curves were assessed by mixed effect linear model using *lmerTest (*v3.1-3). Pairwise statistics were calculated for parametric data by two-tailed t-test and for non-parametric data by Wilcoxon test. A Shapiro test was used to assist in determining if the data were normally distributed.

#### Neoantigen prediction

Preprocessing and variant calling of whole exome sequencing data from Res 499 and B16 cancer cells have been previously described (Benci et al., 2019). Gene expression from paired RNA-seq data was used to filter out variants having transcript read counts of less than three. The MHC-I binding affinities of variants were then predicted using NetMHC version 4.0 for H-2-Kb and H-2-Db using peptide lengths from 8 to 11. High-affinity neoantigens were enriched by removing variants with a predicted MHC-I IC_50_ of greater than 100 nM, and variants with an allelic frequency of 0.5 or greater were also excluded to enrich for variants susceptible to potential LOH. With this filtered list, the genomic contraction of variants in Res 499 versus parental B16 was assessed by examining variants with allele frequencies near-heterozygous (0.2 for a tetraploid genome) in one cell line but subclonal in the other. Significance between the distribution of allelic frequencies between the two groups was estimated by a KS-test.

#### *Aire*-regulated genes

Transcriptional data for mTECs from wild type and *Aire*^-/-^ mice (St-Pierre *et al*., 2015) were obtained from the GEO (GSE65617). Normalized expression values for *Dct* were analyzed using GREIN (https://shiny.ilincs.org/grein).

#### Single cell RNA-seq processing

The *Cell Ranger* pipeline (10x Genomics) was used to process raw data according to the company recommendations (cellranger-mm10-3.0.0). Genes expressed in less than 0.1% cells were filtered out prior to imputation with *SAVER* (v1.1.2). Genes expressed in less than 1% cells and cells with less than 500 detected genes were filtered, and cells with over 10% mitochondrial reads were removed. Variations in UMI count, percent of mitochondrial contamination, and cell cycle were regressed out and samples were integrated using *Seurat* (v.3.1.2). After dimensionality reduction and nearest neighbors clustering, clusters were annotated based on marker gene and differential gene expression.

#### Single cell TCR-seq analysis

The *Cell Ranger* pipeline (10x Genomics) was used to align reads to the mouse genome (*cellranger-vdj-GRCm38-alts-ensemble-2.2.0*). Cells without full-length productive rearrangement were filtered. Clonotypes were defined by unique TCRβ CDR3 sequence; clonotype frequency was calculated within each biological replicate. TCR data was integrated with gene expression data by UMI.

#### Gene set variation analysis

*GSVA* (v.1.32.0) was used to identify enriched gene sets for each CD8 T cell cluster using the per cluster averaged RNA expression data. Gene sets for CD8 T cell subsets were derived using previously reported RNA-sequencing data for CD8 T cell from acute or chronic LCMV infection (Hudson et al., 2019). Gene sets for each subset was extracted by selecting genes with at least a 2-fold increase in each subset compared to all others. For the general exhaustion gene set, TIM3^+^ PD1^+^ CD8 T cells were used as the comparison group. To assess NFAT/AP-1 gene enrichment, *GSVA* was used to determine per cell enrichment of *Fosb*, *Fosl2*, *Fos*, *Junb*, *Jund*, and *Nfatc3* for all CD8 T cells.

### TCR repertoire analysis

Bulk TCR sequencing data from peripheral TRP2 tetramer-positive CD8 T cells from Res 499 tumor bearing mice treated with anti-CTLA4 + anti-PD1 with or without anti-RANKL pre-conditioning were used to characterize features of the TRP2-reactive T cell repertoire. Clonotypes with a frequency of less than 0.01% were excluded. The Jaccard index, Gini coefficient, TCRB V-gene usage, and CDR3 length distribution were determined using the *immunarch* R package. To examine how anti-RANKL impacts the T cell repertoire, the *DeepTCR* Python package (Sidhom *et al*., 2021)was used for deep learning-based supervised repertoire classification. Here, TCRB features that include the CDR3 amino acid sequence, counts, V-genes, and J-genes were used to train the model by 100-fold Monte Carlo cross-validation with the following parameters: *epochs_min* = 10, *size_of_net* = ’small’, *num_concepts* = 64, *hinge_loss_t* = 0.1, *train_loss_min* = 0.1. The learned TCR repertoire features from the model were visualized in low-dimensional space by UMAP using *n* = 50 nearest neighbors and clustered using PhenoGraph, as implemented in the *Rphenograph* R package, with *k* = 200 nearest neighbors. To understand the biochemical properties of the CDR3 amino acid sequences that are representative of each TCR cluster, the CDR3 sequences with a predicted class probability of less than 75% were first filtered. Then, CDR3 sequences from the top clusters exclusive to anti-RANKL + ICB (n = 4), ICB alone (n = 4), or shared between the two treatment groups (n = 6) were extracted and used to construct a consensus amino acid alignment using the *msa* R package. The amino acids from his consensus alignment were then assigned four Atchley factors that quantitates 1) electrostatic charge (EC), 2) molecular size (MS), 3) polarity, accessibility, and hydrophobicity (PAH), and 4) propensity for secondary structure (PSS) using the *HDMD* R package.

In order to predict the probability that a new CD8 T cell clonotype comes from the anti-RANKL TRP2-reactive T cell repertoire, the aforementioned TCRB features (CDR3 amino acid sequence, counts, V-gene, and J-gene) from independent scRNA/TCR-sequencing data of TILs from tumor-bearing mice treated with ICB +/- anti-RANKL was used as test data for the *DeepTCR* model. The RNA expression data was then used to determine the phenotype, *Nur77* expression, and NFAT/AP-1 gene enrichment of the CD8 T cells belonging to each clonotype.

### Human melanoma single-cell RNA sequencing analysis

Single-cell RNA sequencing data from 48 melanoma patients treated with and without ICB (Sade-Feldman et al., 2018) were obtained from the GEO (GSE120575). Data were re-normalized using *sctransform* from the *Seurat* R package (v4.0.1) and denoised using the *SAVER* R package (v1.1.2). After dimensionality reduction and shared nearest-neighbors clustering using *Seurat*, only clusters defined as T cells or innate lymphoid cells were selected and the data re-clustered. T cell and innate lymphoid cell clusters from post-treatment biopsies that predict ICB response was determined using random forest as implemented in the *randomForestSRC* R package. Because responder and non-responder classes were imbalanced, the *imbalanced* function was used with *nodesize* = 1 and *ntree* = 1000. Prediction error rates and variable importance scores were averaged from 100 Monte Carlo simulations. To determine the extent to which predictive CD8 T cell clusters are enriched for genes differentially expressed by mouse CD8 T cell subsets that infiltrate Res 499 tumors, GSVA was performed using upregulated genes with a log fold-change of at least 0.3 from the scRNA-seq mouse CD8 TIL clusters (using *FindAllMarkers* from *Seurat*).

